# The megakaryocytic transcription factor ARID3A suppresses leukemia pathogenesis

**DOI:** 10.1101/2021.04.26.440795

**Authors:** Oriol Alejo-Valle, Karoline Weigert, Raj Bhayadia, Michelle Ng, Stephan Emmrich, Christoph Beyer, Konstantin Schuschel, Christian Ihling, Andrea Sinz, Marius Flasinski, Hasan Issa, Eniko Regenyi, Maurice Labuhn, Dirk Reinhardt, Marie-Laure Yaspo, Dirk Heckl, Jan-Henning Klusmann

**Author notes:** **Corresponding authors**: Jan-Henning Klusmann; Dirk Heckl. Joint last authors (Dirk Heckl, Jan-Henning Klusmann). **Lead contact author**: Jan-Henning Klusmann. **Scientific category:** Hematopoiesis and stem cells.

## Abstract

Given the plasticity of hematopoietic stem/progenitor cells, multiple routes of differentiation must be blocked during acute myeloid leukemia pathogenesis – the molecular basis of which is incompletely understood. Here we report that post-transcriptional repression of transcription factor ARID3A by miR-125b is a key event in megakaryoblastic leukemia (AMKL) pathogenesis. AMKL is frequently associated with trisomy 21 and *GATA1* mutations (GATA1s), and children with Down syndrome are at a high risk of developing this disease. We show that chromosome 21-encoded miR-125b synergizes with *Gata1s* to drive leukemogenesis in this context. Leveraging forward and reverse genetics, we uncover *Arid3a* as the main miR-125b target underlying this synergy. We demonstrate that during normal hematopoiesis this transcription factor promotes megakaryocytic differentiation in concert with GATA1 and mediates TGFβ-induced apoptosis and cell cycle arrest in complex with SMAD2/3. While *Gata1s* mutations perturb erythroid differentiation and induce hyperproliferation of megakaryocytic progenitors, intact ARID3A expression assures their megakaryocytic differentiation and growth restriction. Upon knockdown, these tumor suppressive functions are revoked, causing a dual megakaryocytic/erythroid differentiation blockade and subsequently AMKL. Inversely, restoring *ARID3A* expression relieves the megakaryocytic differentiation arrest in AMKL patient-derived xenografts. This work illustrates how mutations in lineage-determining transcription factors and perturbation of post-transcriptional gene regulation interplay to block multiple routes of hematopoietic differentiation and cause leukemia. Surmounting this differentiation blockade in megakaryoblastic leukemia by restoring the tumor suppressor ARID3A represents a promising strategy for treating this lethal pediatric disease.

**Key points:**

- Repression of megakaryocytic transcription factor *ARID3A* by miR-125b synergizes with *GATA1s* to induce leukemia
- Restoring ARID3A expression relieves megakaryocytic differentiation arrest in megakaryoblastic leukemia

**Graphical abstract:** 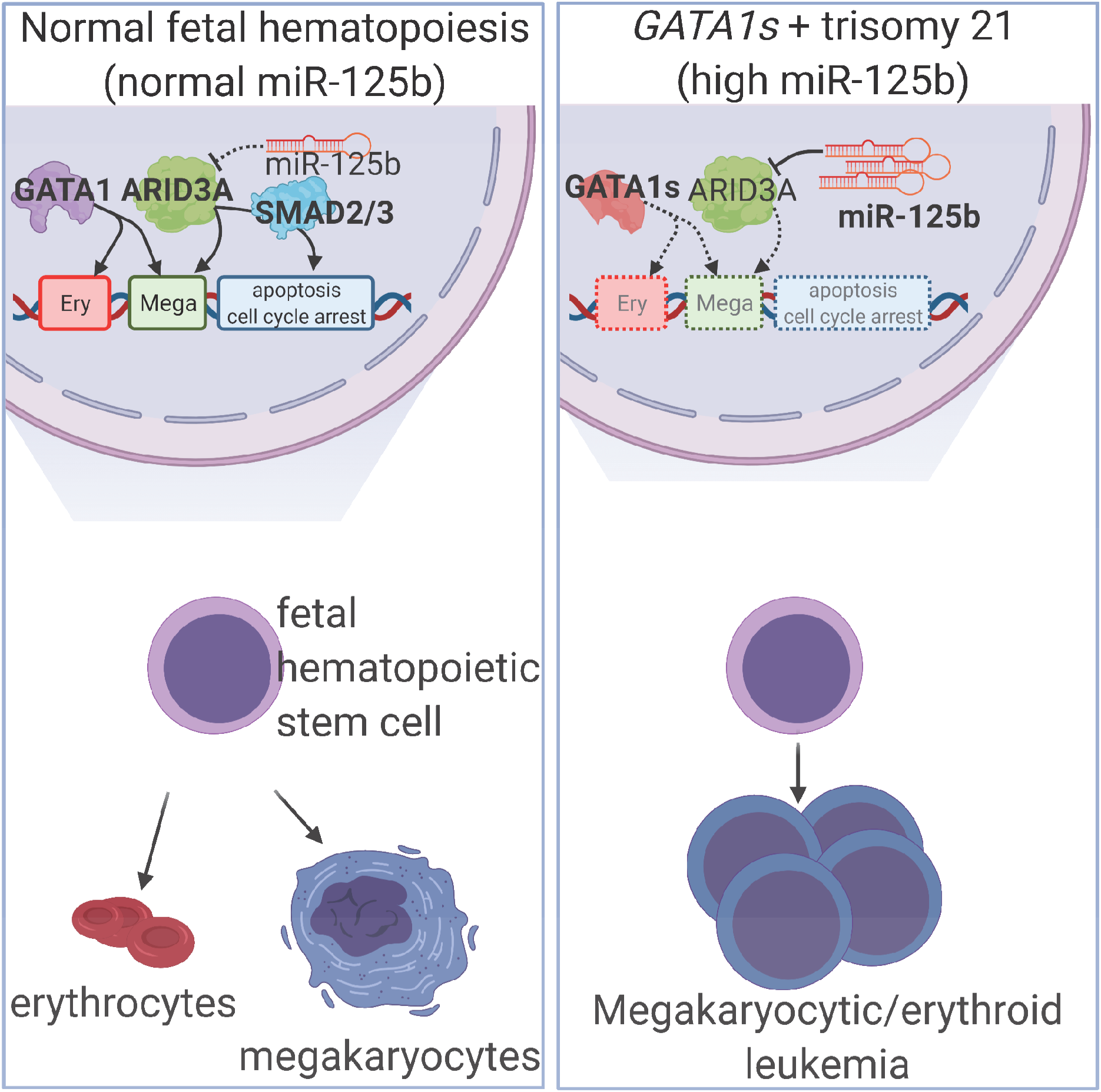

## Introduction

Acute myeloid leukemia (AML) is a hematologic malignancy characterized by the uncontrolled proliferation of immature progenitor cells, which are blocked from differentiating towards normal blood cells. Given the plasticity of hematopoietic stem/progenitor cells (HSPCs), multiple routes of differentiation must be blocked during AML pathogenesis – the molecular basis of which is incompletely understood. In particular, the role of perturbed post-transcriptional gene regulation in leukemic differentiation arrest remains unknown. Understanding how these altered processes lead to leukemic transformation is crucial for developing strategies to overcome the differentiation blockade in AML and for designing related therapeutic concepts, as has been successfully applied in acute promyelocytic leukemia^1^.

Acute megakaryoblastic leukemia (AMKL) is an aggressive subtype of AML. Acquired trisomy 21 is a common feature of AMKL^2^ and children with Down syndrome are at a high risk of developing this disease. Megakaryoblastic leukemia of Down syndrome (ML-DS) is characterized by mutations in the hematopoietic transcription factor GATA1, which occur in fetal hematopoietic stem/progenitor cells (HSPCs) and cause the exclusive expression of a N-terminal truncated protein known as GATA1s – hereafter referred to as *GATA1s* mutations^3–5^. Noteworthy, ML-DS pathogenesis is a stepwise clonal progression evolving from a transient form of leukemia called transient abnormal myelopoiesis (TAM), which occurs in approximately 30% of neonates with Down syndrome. Today it is widely accepted that the interplay between trisomy 21, fetal origin and the mutated GATA1s is necessary and sufficient to cause TAM, as none of these elements lead to a TAM-like phenotype alone^4, 6–8^, yet no additional events/factors are required^9, 10^. During fetal development, GATA1s induces hyperproliferation of human and murine megakaryocytic progenitor cells, but leaves their capacity to differentiate into normal megakaryocytes intact^4, 11^. Meanwhile, *Gata1s* causes anemia in murine fetuses and interferes with normal erythroid differentiation^3, 4^, and *GATA1s* mutations are seen in patients with Diamond-Blackfan anemia^12^. These data suggest that both the erythroid and megakaryocytic differentiation paths are perturbed by the interplay between GATA1s and trisomy 21, and that this combination causes TAM and subsequently ML-DS. Despite the apparent simplicity of this model, the factors on chromosome 21 that underlie this synergy remain enigmatic^*13–19*^. Identifying the oncogenic factors on chromosome 21 will also be pertinent to deciphering its role in perturbed differentiation and leukemogenesis beyond ML-DS^*13*^.

Chromosome 21 harbors the phylogenetically conserved miR-99a~125b cluster, which encodes the miRNAs let-7c, miR-99a and miR-125b^20^. All three are transcribed simultaneously as a polycistron and are highly expressed in TAM, ML-DS and non-DS AMKL^20^. MiR-125b was first described as an oncogenic miRNA in ML-DS^21^ and other types of AML^22, 23^. However, the role of the miRNA tricistron in ML-DS pathogenesis and its function in perturbed differentiation remained an open question. Here we resolved the interplay between *GATA1s*-mutations and the miR-99a~125b cluster and propose a mechanism through which the dual differentiation arrest is achieved in ML-DS and AMKL. By systematically dissecting the miR-125b targetome – the dominant oncogenic member of the cluster in this context – we establish ARID3A as a transcriptional activator that promotes megakaryocytic differentiation in concert with GATA1. These insights will advance our knowledge of the complex regulation of normal hematopoiesis and our understanding of how post-transcriptional regulators of gene expression interact with known oncogenic drivers during leukemic differentiation blockade, as well as during the initiation and progression of cancer.

## Methods

### Reagents and Resources

**Supplemental Table S1** contains a list of all relevant reagents.

### Patient samples

Pediatric AML samples were collected from patients enrolled in AML Berlin-Frankfurt-Münster treatment protocols for children and adolescents. Written informed consent was obtained from all patients and custodians in accordance with the Declaration of Helsinki and local laws and regulations, and the study was approved by the institutional review board of each participating center. For details, see **Supplemental Table S2**.

### Animal studies

All experimental procedures involving mice were performed in accordance with protocols approved by the local authorities (Landesverwaltungsamt Niedersachsen/Sachsen-Anhalt). B6J.129(Cg)-Gt(ROSA)26Sor^tm1.1(CAG-cas9*,-EGFP)Fezh^/J) (Jackson Laboratory, RRID: IMSR_JAX:026179), C57BL/6J mice (Charles River, RRID: IMSR_JAX:000664) and C;129S4-*Rag2^tm1.1Flv^Csf1^tm1(CSF1)Flv^ Csf2*/*Il3^tm1.1(CSF2,IL3)Flv^Thpo^tm1.1(TPO)Flv^Il2rg^tm1.1Flv^*Tg(SIRPA)1Flv/J (MISTRG) mice (Regeneron Pharmaceuticals)^24^ were maintained in a specific pathogen free environment in individual ventilated cages and fed with autoclaved food and water at Martin-Luther-University Halle-Wittenberg. Syngeneic transplantation and patient-derived xenograft protocols have been described previously^9, 25^.

### Statistical analysis

All statistical analyses were carried out using GraphPad Prism 8. Data were analyzed by Student t test (two-tailed). Survival was measured according to the Kaplan–Meier method and analyzed by log-rank (Mantel–Cox) test. P<0.05 was considered significant. All statistical tests and sample numbers are disclosed in the respective figure legends/Supplemental tables.

### Data availability

RNA-Seq gene expression data have been deposited in NCBI’s Gene Expression Omnibus and are accessible through GEO Series accession numbers: GSE169738, GSE169739 and GSE169740. Data regarding shRNA-positive screen is accessible through European Genome-phenome Archive accession number: PRJEB43922. Mass spectrometry proteomics data have been deposited to the ProteomeXchange Consortium via the PRIDE partner repository with the dataset identifier PXD025027. Other remaining data are available within the Article and Supplemental Files, or from the authors upon request.

Additional and detailed methods can be found in **Supplemental Methods.**

## Results

### miR-125b cooperates with *Gata1s* to induce megakaryoblastic leukemia

We previously generated a murine model of *Gata1s*-driven preleukemia, in which we introduce the *Gata1s* mutation in the fetal context using the CRISPR-Cas9 system^9^. The generated *Gata1s* fetal liver cells (FLCs) show a hyperproliferative phenotype but unperturbed terminal megakaryocytic differentiation *in vitro*^4^. Importantly, upon transplantation into syngeneic recipients (C57Bl/6J), *Gata1s*-FLCs become abundant in the peripheral blood and then disappear over time^9^. Thus, our murine preleukemic model constitutes a suitable platform to study and identify *GATA1s*-cooperating events, and we leveraged this system in combination with fluorescence-based lentiviral barcoding to dissect the role of miR-99a, let-7c and miR-125b – the members of the miR-99a~125b tricistron – in TAM/ML-DS pathogenesis. Each of the miRNAs was associated with a reporter fluorescent protein (dTomato, GFP and mTagBFP2, respectively), which enabled monitoring and competition between all possible miRNA permutations (**Figure 1A** and **Supplemental Figure 1A**). This experimental setup revealed major synergy between *Gata1s* and miR-125b – but no other miRNA alone or in combination leading to a 131-fold expansion of FLCs (**Figure 1A** and **Supplemental Figure 1B-C**). MiR-125b further enhanced the *Gata1s*-induced expansion of CD117^+^ stem/progenitor cells, including CD117^+^CD41^+^ megakaryocytic progenitors (**Figure 1B** and **Supplemental Figure 1D**). Correspondingly, the fraction of mature CD41^+^CD42d^+^ megakaryocytes was reduced, implying perturbed megakaryocytic differentiation, while erythroid differentiation was enhanced (**Figure 1B**). In methylcellulose-based colony forming unit (CFU) assays with thrombopoietin (Thpo) as the only cytokine, *Gata1s*-miR-125b FLCs were able to generate megakaryocyte-like CFUs with replating potential, in contrast to *Gata1s* control cells (**Figure 1C-D**). Complete cytokine conditions led to the formation of fewer but larger CFUs – including colonies with blast-like features – in *Gata1s-*miR-125b FLCs (**Figure 1C and Supplemental Figure 1E-F**), with a relative increase of erythroid colonies (CFU-E/ BFU-E; **Figure 1E**).

**Figure 1.**
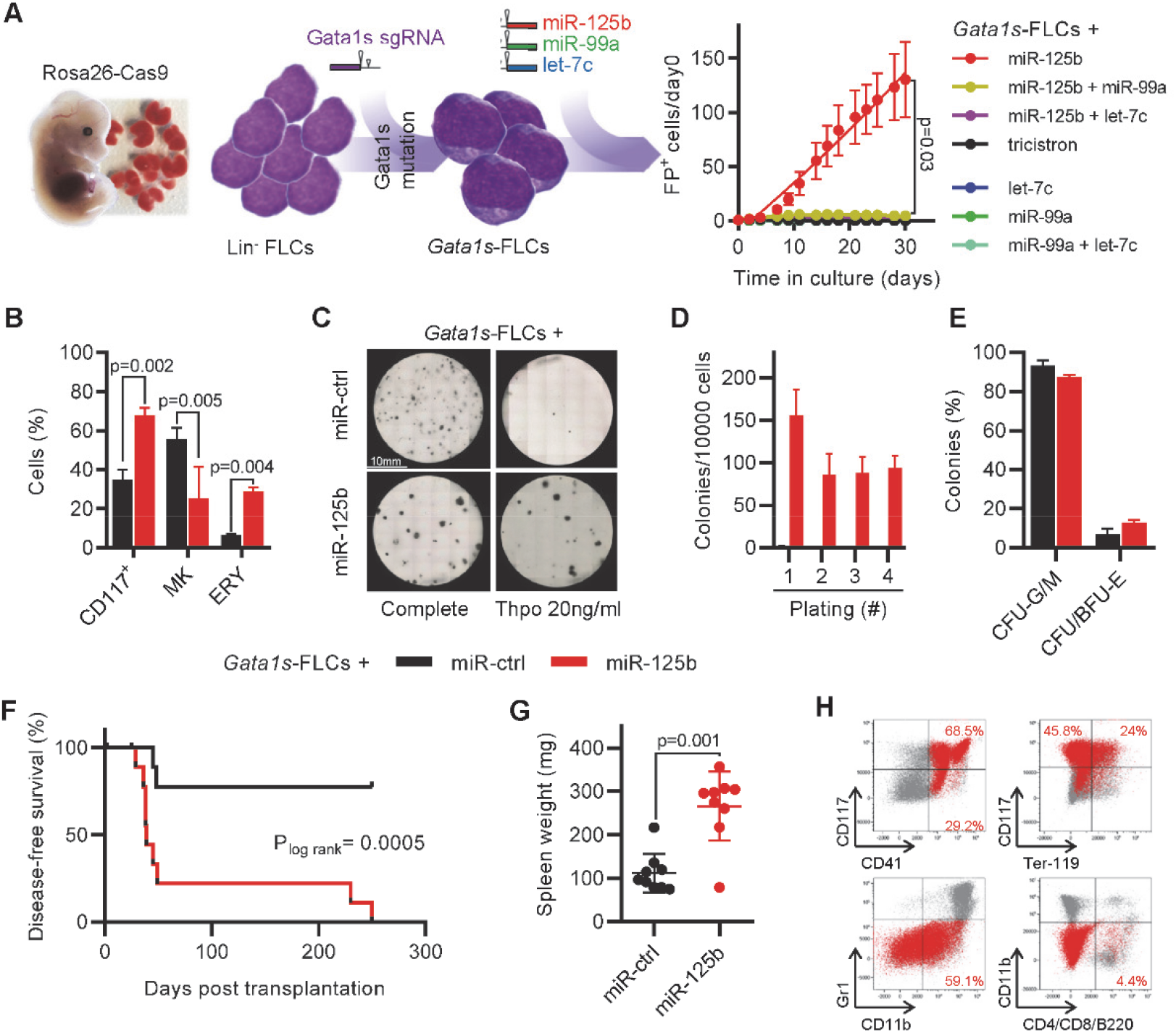
miR-125b cooperates with *Gata1s* to induce leukemia *in vivo.* (A) Schematic of *in vitro* and *in vivo* setup for modeling synergy between Gata1s and the members of miR-99a~125b tricistrons (left). Percentage of *Gata1s*-FLCs transduced with different miRNA permutations (marked by dTomato [miR-125b], mTagBFP2 [let-7c] and GFP [miR-99a]) normalized to day 0 (n=4, paired t-test) (right). (B) Bar graph showing the percentage of stem/progenitor cells (CD117^+^), mature megakaryocytes (MK, CD41^+^CD42d^+^) and erythroid cells (ERY, CD71^+^Ter-119^+^) after 6 days of differentiation. Cells shown are *Gata1s*-FLCs transduced with miR-125b or miR-ctrl (n=3, paired t-test). (C) Representative methylcellulose-based CFU assays from one of n=4 independent experiments. Depicted are *Gata1s-FLC*s transduced with miR-ctrl or miR-125b in complete or low (Thpo 20ng/mL) cytokine conditions. (D) Bar graph showing the number of megakaryocytic colonies after serial replating of miR-125b *Gata1s*-FLCs in methylcellulose-based media with low (Thpo 20ng/mL) cytokine conditions compared to miR-ctrl *Gata1s*-FLCs. (E) Classification of colonies after plating miR-125b or miR-ctrl *Gata1s*-FLCs in methylcellulose-based CFU assays under complete cytokine conditions (paired t-test). CFU-G/M: granulocytic (CFU-G), monocytic (CFU-M) and granulocytic/monocytic (CFU-GM); CFU/BFU-E: erythroid. (F-H) Kaplan-Meier survival curve of recipients transplanted with *Gata1s*-FLCs overexpressing miR-125b or miR-ctrl (n=10, log-rank test) (F), spleen weight (unpaired t-test) (G), and representative flow cytometry plots of bone marrow (BM)-derived leukemic cells in the diseased mice (H). *Gata1s*-FLC-derived blasts overexpressing miR-125b are highlighted in red. Average percentage of miR-125b^+^ blasts belonging to each immunophenotype are indicated in the corresponding gate (H). Data are presented as mean ± SD.

Most importantly, upon transplantation into syngeneic C57BL6/J recipients, miR-125b-transduced *Gata1s*-FLCs caused high penetrance (90%) megakaryoblastic leukemia after a latency of 39 days (median disease-free survival, **Figure 1F-H**). Leukemic blasts engrafted and induced leukemia in secondary recipients with 100% penetrance, underlining the leukemic potential of the *Gata1s*-miR-125b combination (**Supplemental Figure 1G-H**).

These results from the murine *Gata1s* preleukemic model concur with our previous study^21^, and confirm miR-125b as the sole driver of the miR-99a~125b tricistron’s synergy with *Gata1s* in FLCs and the ensuing ML-DS-like megakaryocytic leukemia *in vivo*.

### shRNA-based positive selection screening in combination with RNA-sequencing identifies miR-125b targets that synergize with *Gata1s*

To identify downstream targets of miR-125b that mediate its synergy with *Gata1s*, we performed an *in vitro* short hairpin RNA (shRNA)-based positive selection screen individually probing 220 predicted miR-125b targets that are downregulated by miR-125b in HSPCs (**Figure 2A** and **Supplemental Table 3**)^20, 26^. Next generation sequencing (NGS)-based shRNA quantification 4 days and 30 days post-transduction followed by MAGeCK analysis^27^ revealed a continuous significant enrichment of shRNAs targeting 13 candidate genes (**Figure 2A**, **Supplemental Figure 2A** and **Supplemental Table 4**). We complemented the data obtained from the shRNA screening with gene expression profiling (RNA-seq) after switching miR-125b expression on and off using a doxycycline-inducible lentiviral expression system (**Figure 2B**). We showed that sustained miR-125b expression, e.g. continuous addition of doxycycline, is required for the enhanced proliferation of *Gata1s*-FLCs and related transcriptional changes, such as the induction of ML-DS and stem cell gene expression signatures^28^ as well as of MYC target genes^29^ (**Figure 2B**, **Supplemental Figure 2B-C** and **Supplemental Table 5**). Hence, candidate targets of miR-125b should be downregulated upon miR-125b induction (doxycycline addition) and become re-expressed upon miR-125b release (doxycycline withdrawal). Only one candidate followed this expression pattern, in addition to being significantly enriched in the shRNA screen: the transcription factor *Arid3a* (**Figure 2C**). This convergence from two complementary approaches implicates *Arida3* as the primary player in the synergy between miR-125b with *Gata1s*.

**Figure 2.**
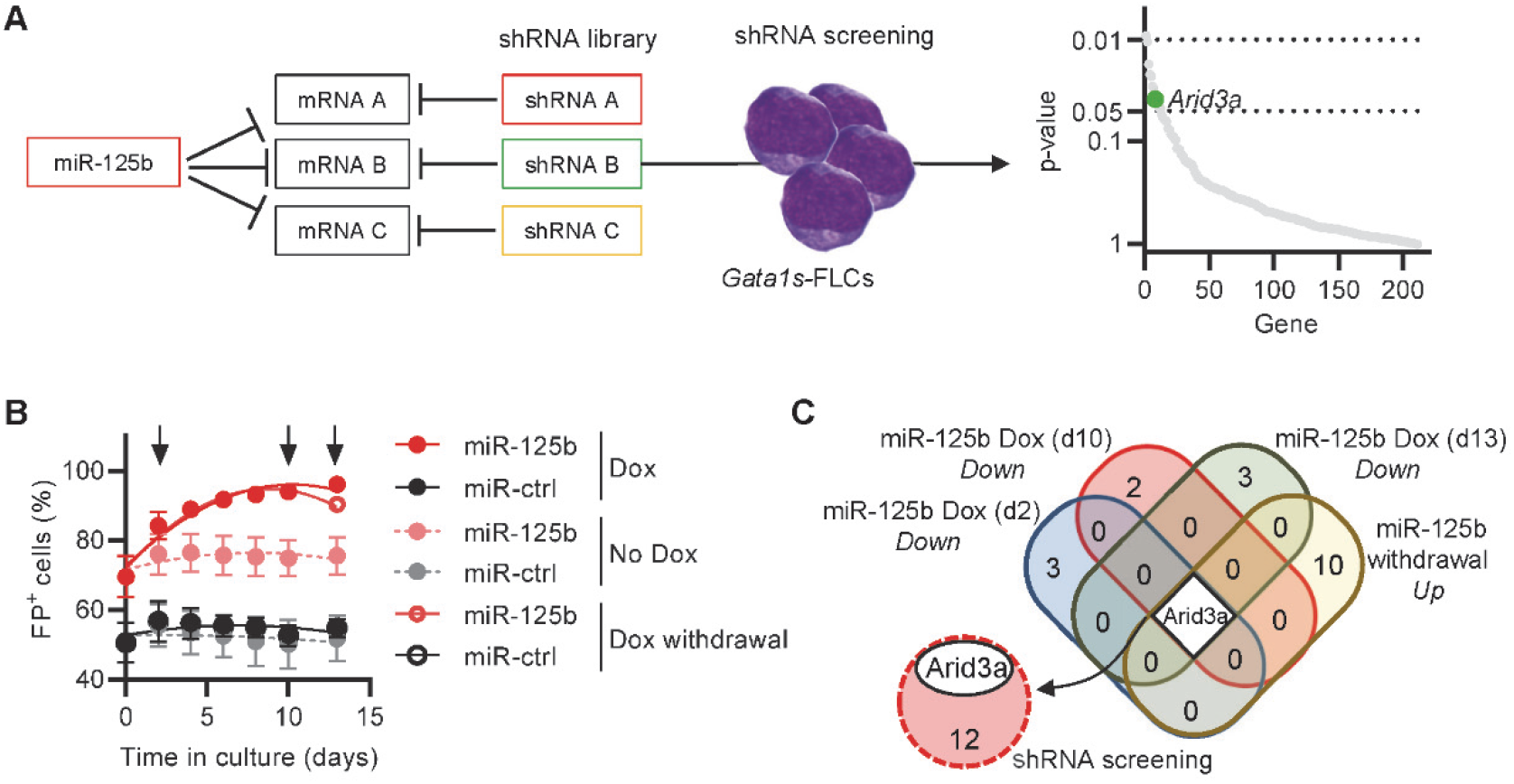
shRNA-based positive selection screening in combination with RNA-sequencing identifies miR-125b targets that synergize with Gata1s. (A) Schematic of shRNA positive selection screen, where a pool of shRNAs directed against miR-125b target genes mimics the effect of miR-125b (left). Dot plot showing significantly enriched shRNA-targeted genes (right). *Arid3a* highlighted in green. (B) Percentage of *Gata1s*-FLCs expressing doxycycline-regulated miR-125b or miR-ctrl upon addition or removal of doxycycline (500ng/mL; n=4). Data presented as mean ± SD. (C) Venn diagram of differentially expressed genes upon miR-125b modulation, overlapped with the results of the shRNA positive selection screen (from (A)).

### *Arid3a* knockdown mimics the miR-125b phenotype in *Gata1s*-FLCs

Next, we sought to validate *Arid3a* as the primary target of miR-125b in this context. We confirmed repression of ARID3A protein as well as direct targeting of the *Arid3a*/*ARID3A* 3’UTR by miR-125b (**Supplemental Figure 3A-D**). Knockdown of *Arid3a* in *Gata1s*-FLCs using two efficient shRNAs (shArid3a #2 and shArid3a #3, **Supplemental Figure 3A** and **Supplemental Table 6**) mimicked the miR-125b pro-proliferative phenotype and megakaryocytic differentiation arrest (**Figure 3A-B**). In CFU assays (Thpo only), *Arid3a* knockdown resulted in the formation of megakaryocyte-like CFUs with high replating potential (**Figure 3C-D** and **Supplemental Figure 3E**), while in complete cytokine conditions, knockdown of *Arid3a* led to a relative increase in erythroid colonies (CFU-E/ BFU-E) and colony size (**Supplemental Figure 3F-H**). Inversely, restoring *Arid3a* levels in *Gata1s*-miR-125b FLCs via cDNA overexpression reestablished megakaryocytic differentiation while impeding miR-125b-driven proliferation (**Figure 3E-F**) and CFU formation (Thpo only, **Figure 3G-H**). Most importantly, knockdown of *Arid3a* in *Gata1s*-FLCs was sufficient to recapitulate the miR-125b-induced leukemic phenotype *in vivo*, leading to the development of CD117^+^CD41^+^ megakaryoblastic leukemia with a short latency of 71 days (median disease-free survival, p=0.02), accompanied by marked splenomegaly (**Figure 3I-K**).

**Figure 3.**
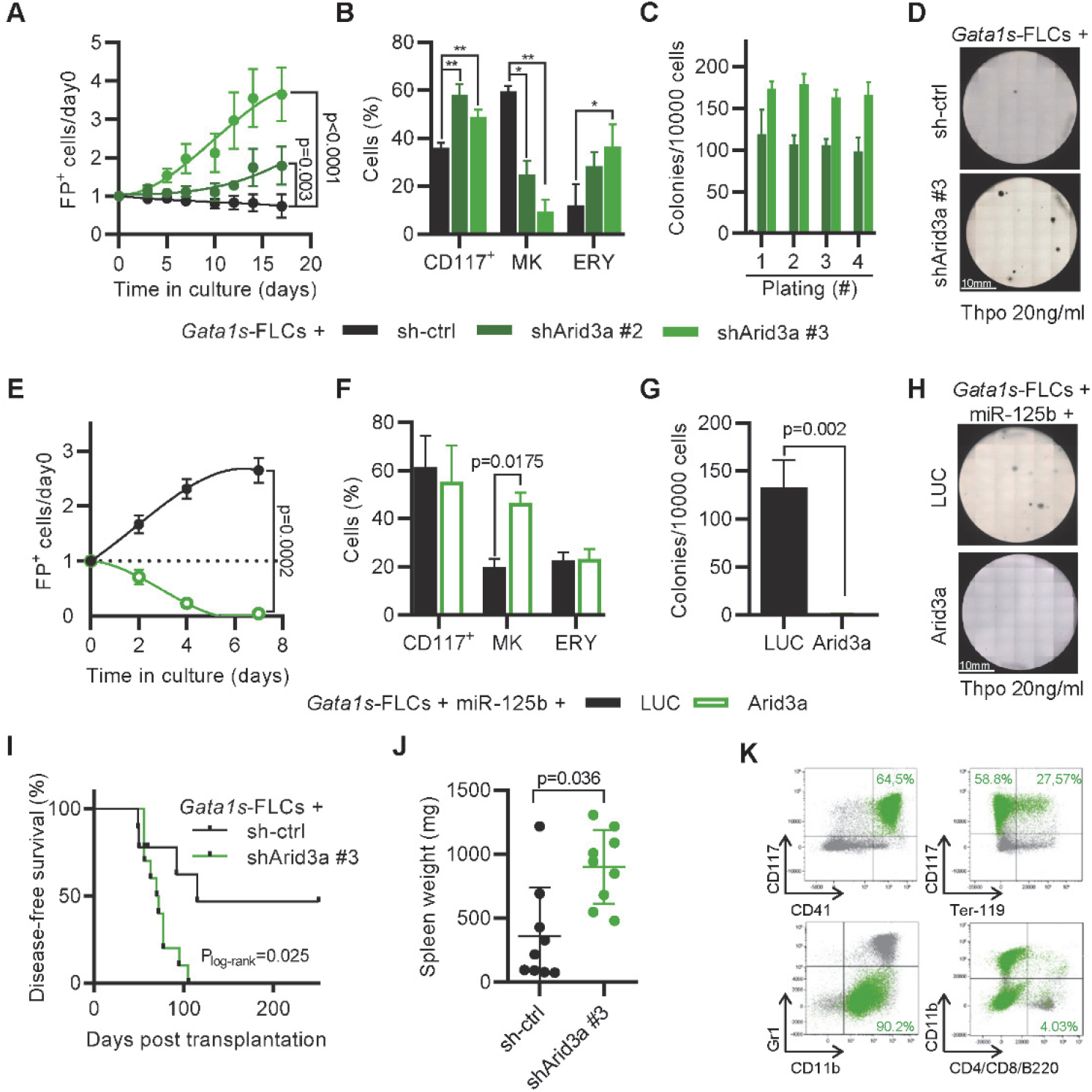
*Arid3a* knockdown mimics the miR-125b phenotype in *Gata1s*-FLCs. (A) Percentage of transduced (sh-ctrl or shArid3a #2 or #3) *Gata1s*-FLCs compared to day 0 (n=4, paired t-test). (B) Bar graph showing the percentage of stem/progenitor cells (CD117^+^), mature megakaryocytes (MK, CD41^+^ CD42d^+^) and erythroid cells (ERY, CD71^+^Ter-119^+^) in transduced (sh-ctrl (black) or shArid3a #2 or #3 (dark or light green, respectively)) *Gata1s*-FLC after 6 days of differentiation of. *= p<0.05; ** = p<0.01; n=3 (paired t-test). (C-D) Bar graph showing the number of megakaryocytic colonies after serial replating of transduced (sh-ctrl (black) or shArid3a #2 or #3 (dark or light green, respectively)) *Gata1s*-FLCs in methylcellulose-based media with low concentrations of Thpo (20ng/mL); serial replating number given on x-Axis (C). Representative image of one of n=3 independent experiments (D). (E) Percentage of transduced (*Arid3a* (green) or LUC cDNA (black)) miR-125b-*Gata1s*-FLCs normalized to day 0 (n=4, paired t-test). (F) Percentage of stem/progenitor cells (CD117^+^), mature megakaryocytes (MK, CD41^+^CD42d^+^) and erythroid cells (ERY, CD71^+^Ter-119^+^) in transduced (*Arid3a* (green) or LUC cDNA (black)) miR-125b-*Gata1s*-FLCs after 6 days of differentiation (n=3, paired t-test). (G and H) Bar graph depicting the number of colonies after plating transduced (*Arid3a* or LUC cDNA) miR-125b *Gata1s*-FLCs in methylcellulose-based media with low Thpo conditions (20ng/mL) (paired t-test) (G). Representative image of one of n=3 independent experiments (H). (I-K) Kaplan-Meier survival curve of C57BL-6J recipients upon transplantation of transduced (sh-ctrl or shArid3a) *Gata1s*-FLCs (log-rank test) (I), spleen weights (unpaired t-test) (J), and representative flow cytometry plots of bone marrow (BM)-derived leukemic cells from the diseased mice. shArid3a-expressing *Gata1s*-FLC-derived blasts are highlighted in green. Average percentage of shArid3a^+^ blasts belonging to each immunophenotype is indicated in the corresponding gate (K). All data are presented as mean ± SD.

In summary, these results strongly implicate *Arid3a* as the main target of miR-125b that drives the synergy with *Gata1s* in leukemogenesis *in vitro* and *in vivo*.

### ARID3A promotes terminal megakaryocytic differentiation

ARID3A has been extensively studied in the B-cell lineage – also in the context of miR-125b^30^ – where it upregulates IgH transcription in activated B-cells^31^. More recent reports have highlighted its role in myelopoiesis and erythropoiesis^32, 33^. Gene expression analysis in stringently sorted blood cells (R.B. and J.-H.K, unpublished data) showed that *ARID3A* expression is elevated in megakaryocytes compared to erythroid cells (**Figure 4A**), indicating that ARID3A also acts as a regulator of megakaryopoiesis. Accordingly, shRNA-mediated knockdown of *Arid3a* greatly enhanced the colony-forming and replating capacity of wild type (WT) FLCs with Thpo as the only cytokine and impaired terminal megakaryocytic differentiation in the liquid culture (**Figure 4B-C**). Inversely, *Arid3a* overexpression promoted megakaryocytic differentiation at the expense of erythroid differentiation (**Figure 4D**), which was also evident in CFU assays **(Figure 4E** and **Supplemental Figure 4A)**. To verify the effects of *Arid3a* on megakaryopoiesis and erythropoiesis *in vivo*, we transplanted shRNA-transduced FLCs into lethally irradiated syngeneic recipients. After 16 weeks, we observed a marked increase in the percentage of CD71^+^Ter-119^+^ early and CD71^−^Ter-119^+^ late erythroblasts^34^ as well as of CD41^+^CD42d^−^ immature megakaryocytic cells in the bone marrow of the recipient mice upon *Arid3a* knockdown, while the percentage of CD41^+^CD42d^+^ mature megakaryocytes was not significantly changed (**Figure 4F**). Other lineages were not affected (**Supplemental Figure 4B**).

**Figure 4.**
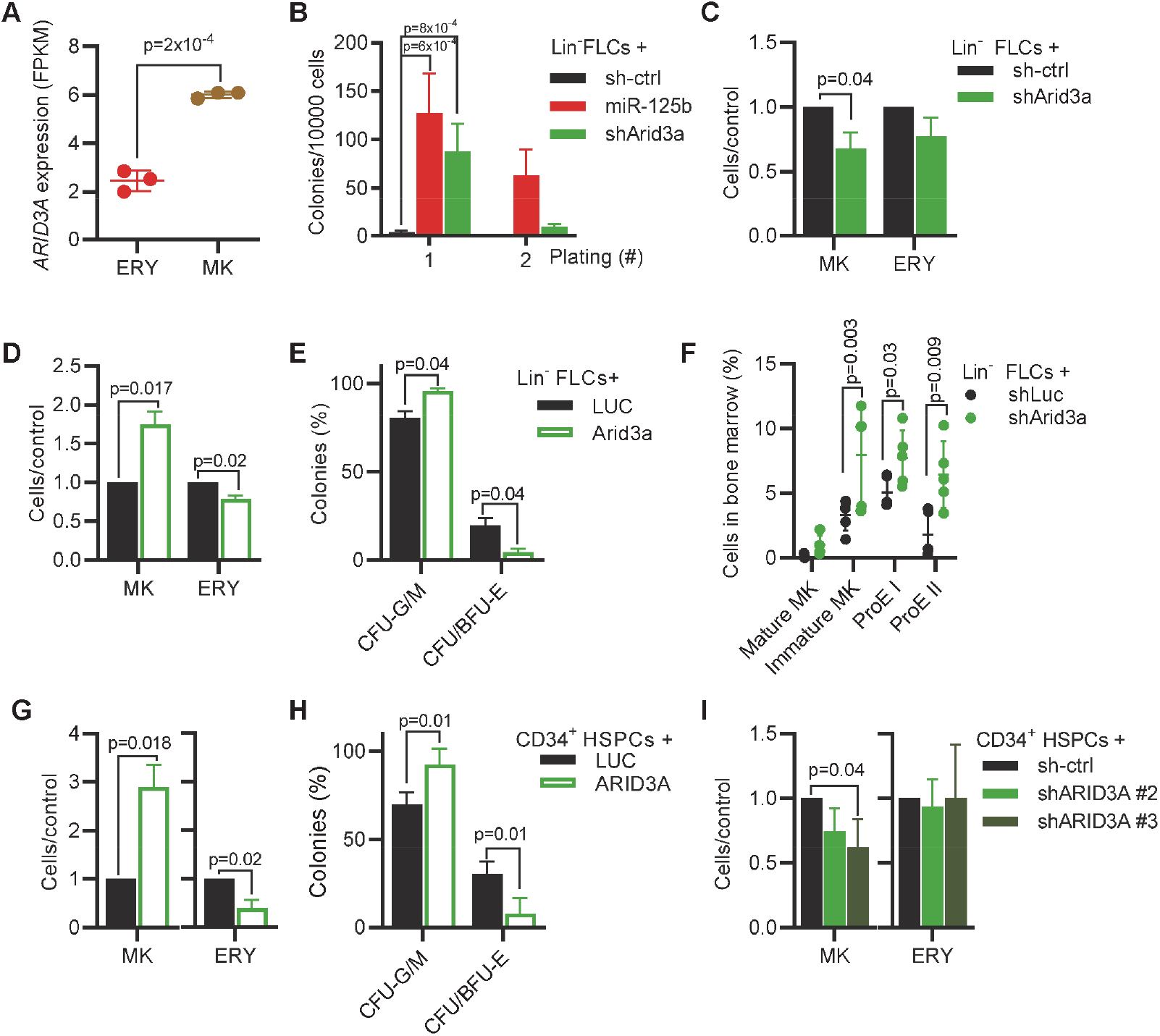
ARID3A promotes terminal megakaryocytic differentiation. (A) *ARID3A* expression (FPKM) in megakaryocytic (CD41^+^CD61^+^) and erythroid cells (CD71^+^) derived from human CD34^+^ HSPCs after 7 days of differentiation in megakaryocytic- or erythroid-promoting culture conditions, respectively (unpaired t-test). (B) Bar graph showing the number of colonies after replating transduced (sh-ctrl, miR-125b or shArid3a) murine FLCs in low Thpo methylcellulose-based CFU-assays (n=6, paired t-test). Serial replating number given on x-Axis. (C-D) Ratio of terminal megakaryocytic (MK, CD41^+^CD42d^+^) and erythroid cells (ERY, CD71^+^Ter-119^+^) in shArid3a-versus sh-ctrl-transduced murine FLCs (C) and *Arid3a*-versus LUC-transduced FLCs (D) after 6 and 4 days of differentiation, respectively (n=3, paired t-test). (E) Classification of colonies after plating transduced (*Arid3a* or LUC cDNA) murine FLCs in methylcellulose-based CFU assays under complete cytokine conditions (n=3, paired t-test). CFU-G/M: granulocytic (CFU-G), monocytic (CFU-M) and granulocytic/monocytic (CFU-GFM); CFU/BFU-E: erythroid. (F) Bar graph showing the percentage of mature megakaryocytes (Mature MK, CD41^+^CD42d^+^), immature megakaryocytes (Immature MK, CD41^+^CD42^−^), early erythroblasts (ProE I, CD71^+^Ter-119^+^) and late erythroblasts (ProE II, CD71^−^Ter-119^+^) in the BM of mice transplanted with Lin^−^ FLCs transduced with sh-ctrl (black) or shArid3a (green). shRNA^+^ cells shown. n=5 (unpaired t-test). (G) Percentage of differentiated cells after transducing human CD34^+^ HSPCs with *ARID3a* cDNA, normalized to HSPCs transduced with LUC. (Left) Percentage of mature megakaryocytes (CD41^+^/CD61^+^/CD42^+^) after 11 days of differentiation in media promoting megakaryocytic differentiation. (Right) Percentage of mature erythroid cells (CD71^+^CD235a^+^) after 7 days of differentiation in media promoting erythroid differentiation. (n=6, paired t-test). (H) Classification of colonies after plating *ARID3A*- or LUC-expressing human CD34^+^ HSPCs in methylcellulose-based CFU assays under complete cytokine conditions (n=6, unpaired t-test). CFU-G/M: granulocytic (CFU-G), monocytic (CFU-M) and granulocytic/monocytic (CFU-GFM); CFU/BFU-E: erythroid. (I) Percentage of terminally differentiated cells after transducing human CD34^+^ HSPCs with shRNAs targeting *ARID3A*, normalized to sh-ctrl. (Left) Percentage of mature megakaryocytes (CD41^+^CD61^+^CD42^+^) after 11 days of differentiation in media promoting megakaryocytic differentiation. (Right) Percentage of mature erythroid cells (CD71^+^CD235a^+^) after 7 days of differentiation in media promoting erythroid differentiation (n=3, paired t-test). Data are presented as mean ± SD.

To verify the relevance of our findings to human hematopoiesis, we tested human CD34^+^ HSPCs. Under conditions promoting megakaryocytic differentiation, *ARID3A* overexpression strongly promoted terminal differentiation, as determined by the presence of CD41^+^CD61^+^CD42b^+^ cells (**Figure 4G**). Inversely, under conditions promoting erythroid differentiation, *ARID3A* caused a 60% reduction in mature CD71^+^CD235a^+^ erythroid cells (**Figure 4G**), and in CFU assays, we observed a decrease in CFU/BFU-E (**Figure 4H** and **Supplemental Figure 4C**). In contrast, shRNA-mediated knockdown of *ARID3A* impaired megakaryocytic differentiation (**Figure 4I**, **Supplemental Figure 4D** and **Supplemental Table 6**).

These data reveal a previously unknown role for ARID3A in the regulation of megakaryocytic differentiation, and confirm previous reports suggesting its negative impact on erythroid differentiation^32^.

### ARID3A acts in concert with GATA1 to activate megakaryocytic transcriptional programs

To define the function of ARID3A during megakaryopoiesis and leukemogenesis, we proceeded with the molecular characterization of this transcription factor. To elucidate whether the observed effects were mediated by its activating or repressing function^35, 36^, we fused the KRAB inhibitor domain^37^ and the VP64 activator^38^ to the N-terminus of ARID3A (**Supplemental Figure 5A**). Lentiviral *Arid3a* expression reduced the proliferation of *Gata1s*-FLCs cells – an effect that was further enhanced by the addition of VP64 and abrogated by the addition of the KRAB (**Supplemental Figure 5B**), suggesting that ARID3A acts mainly through transcriptional activation in this context. These data were supported by gene expression profiling on pre-leukemic *Gata1s*-FLCs, as well as on *Gata1s*-FLCs after doxycycline-induced knockdown (shArid3a) or overexpression of *Arid3a* (**Figure 5A** and **Supplemental Figure 5C-E**). Differentially expressed genes occupied by ARID3A^39^ were mostly upregulated upon *Arid3a* overexpression and downregulated upon *Arid3a* knockdown (89% and 63.4% respectively), including several megakaryocytic genes such as *Meis1* and *Slc2*a3^40, 41^ (**Supplemental Figure 5F-H**).

**Figure 5.**
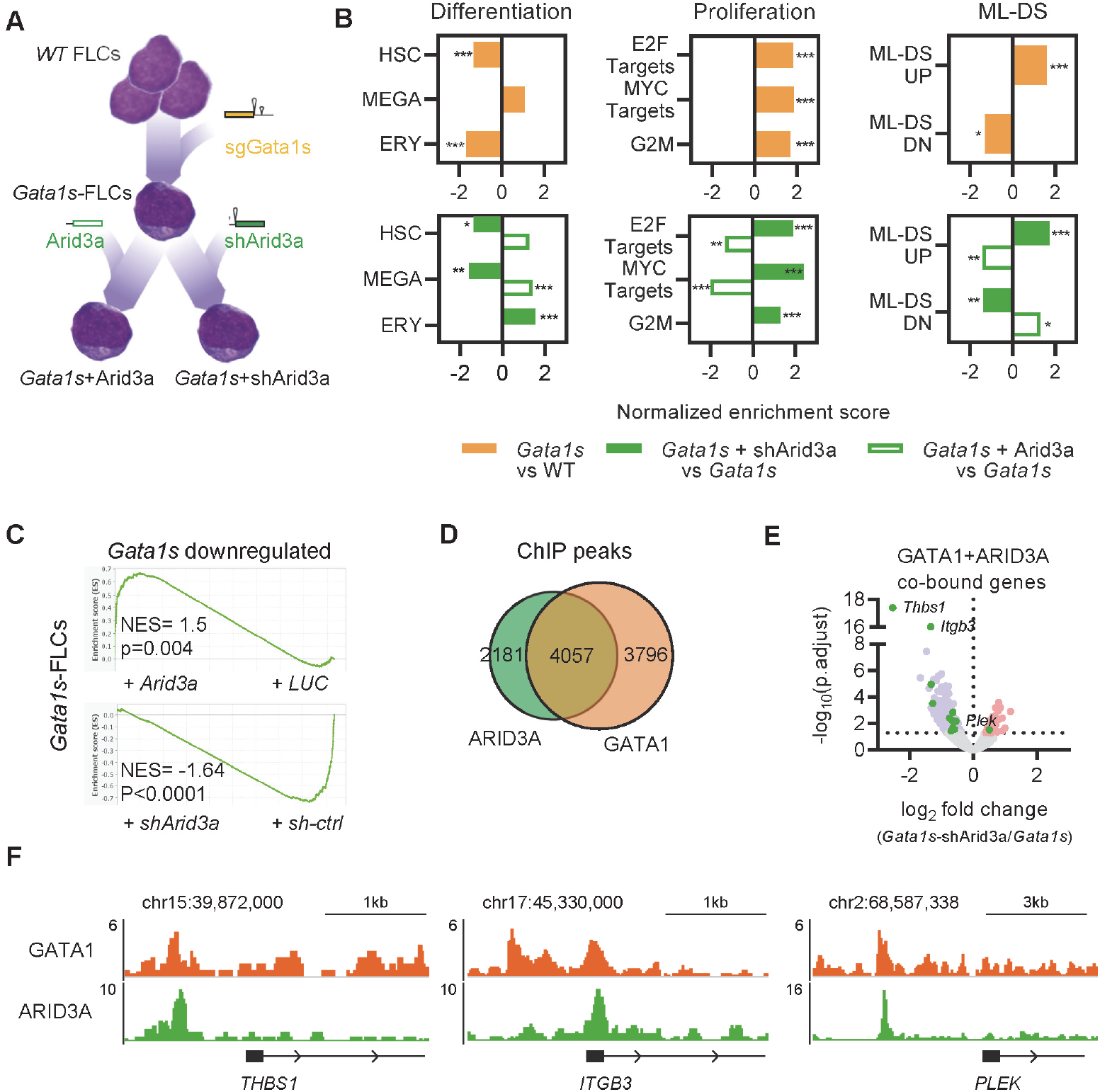
ARID3A acts in concert with GATA1 to activate megakaryocytic transcriptional programs. (A) Modelling the molecular mechanism of TAM to ML-DS transition via sequential *Gata1s* and Arid3a-repression: *Gata1s*-FLCs were expanded for three weeks and then transduced with either *Arid3a* cDNA, shArid3a or their respective controls. RNA samples were obtained from each of the 4 different conditions and subjected to RNA-Sequencing-based gene expression analysis. (B) Bar graphs showing normalized enrichment scores from up- or downregulated gene sets involved in hematopoietic differentiation, cell proliferation and ML-DS progression. *Gata1s*-FLCs were compared against WT FLCs (top); *Arid3a* and shArid3a *Gata1s*-FLCs were compared against their respective *Gata1s*-FLCs controls (LUC or sh-ctrl) after doxycycline induction. *= p<0.05; ** = p<0.01; ***=p<0.001. (C) GSEA enrichment plots showing genes downregulated by the *Gata1s* mutation in FLCs, and their response to *Arid3a* modulation in *Gata1s*-FLCs. (D) Venn diagram of ARID3A and GATA1 ChIP-seq peaks in K562 cells from the ENCODE datasets ^39^. (E) Volcano plot showing differential expression of genes bound by both ARID3A and GATA1 in shArid3a *Gata1s*-FLCs. Genes involved in megakaryocytic differentiation are highlighted in green; significantly downregulated and upregulated genes in blue and red, respectively. (F) IGV snapshots of megakaryocytic genes co-regulated by ARID3A and GATA1 (K562 cells, ENCODE datasets). Scale (y-axis) represents fold enrichment of normalized reads compared to the IgG control. Scale and chromosome location are shown (top).

Next, we determined in FLCs the gene expression changes induced by the *Gata1s* mutation and the additive effects of *Arid3a* repression. As previously described^11, 42^, gene set enrichment analysis (GSEA)^43^ revealed a marked reduction of erythroid genes and concurrent activation of pro-proliferative genes, including MYC and E2F targets, induced by the *Gata1s* mutation (**Figure 5A-B** and **Supplemental Table 7**). On this background, knockdown of *Arid3a* impaired the expression of megakaryocytic genes, which was accompanied by a further amplification of pro-proliferative gene expression and induction of a ML-DS gene expression signature, whereas the opposite was true upon *Arid3a* overexpression (**Figure 5B** and **Supplemental Table 7**). We note that these gene expression changes were highly reminiscent of miR125b overexpression (**Supplemental Figure 2B**).

Interestingly, GSEA analysis also indicated that *Arid3a* knockdown enhances the *Gata1s* gene expression signature. Genes that were downregulated upon loss of full-length GATA1 due to the *Gata1s* mutation were further repressed by *Arid3a* knockdown and re-induced by *Arid3a* overexpression (**Figure 5C**), suggesting that ARID3A maintains expression of these genes at the preleukemic stage. In accordance with this, two-thirds of ARID3A-occupied genes are also bound by GATA1^39^, including several genes involved in megakaryocytic differentiation such as *THBS1*, *ITGB3* (CD61) and *PLEK*^44–46^ (**Figure 5D-F**). Importantly, these genes show similar expression patterns in miR-125b- and shArid3a-transduced *Gata1*s*-*FLCs (**Supplemental Figure 5G**), but do not possess miR-125b binding sites in their 3’UTRs as assessed by TargetScan^26^, suggesting that they are indirectly regulated via miR-125b-mediated knockdown of *Arid3a*. Co-immunoprecipitation (Co-IP) followed by western blot excluded a direct interaction between ARID3A and GATA1 (data not shown), rather suggesting a functional interaction at the promoter level.

Altogether, these data establish ARID3A as a transcriptional activator that promotes megakaryocytic differentiation in concert with GATA1 and that its repression is a key event in megakaryoblastic leukemia (AMKL) pathogenesis.

### ARID3A interacts with SMAD2/3 and promotes TGFβ pathway activation

To further elucidate the tumor suppressive function of ARID3A in the context of ML-DS, we probed the ARID3A protein interaction network through Co-IP of ARID3A in the ML-DS cell line CMK, followed by liquid chromatography with tandem mass spectrometry (LC-MS/MS) (**Figure 6A**). Thirty-seven proteins were significantly enriched, including previously described ARID3A interactors such as ARID3B and PML (**Figure 6B** and **Supplemental Table 8**)^47, 48^. Interestingly, one of the top interaction partners was SMAD2, a downstream effector of the TGFβ pathway. To confirm the LC-MS/MS results, we performed western blots on ARID3A Co-IP samples, and showed a direct interaction between ARID3A and SMAD2 or its paralog SMAD3^49^ (**Figure 6C**).

**Figure 6.**
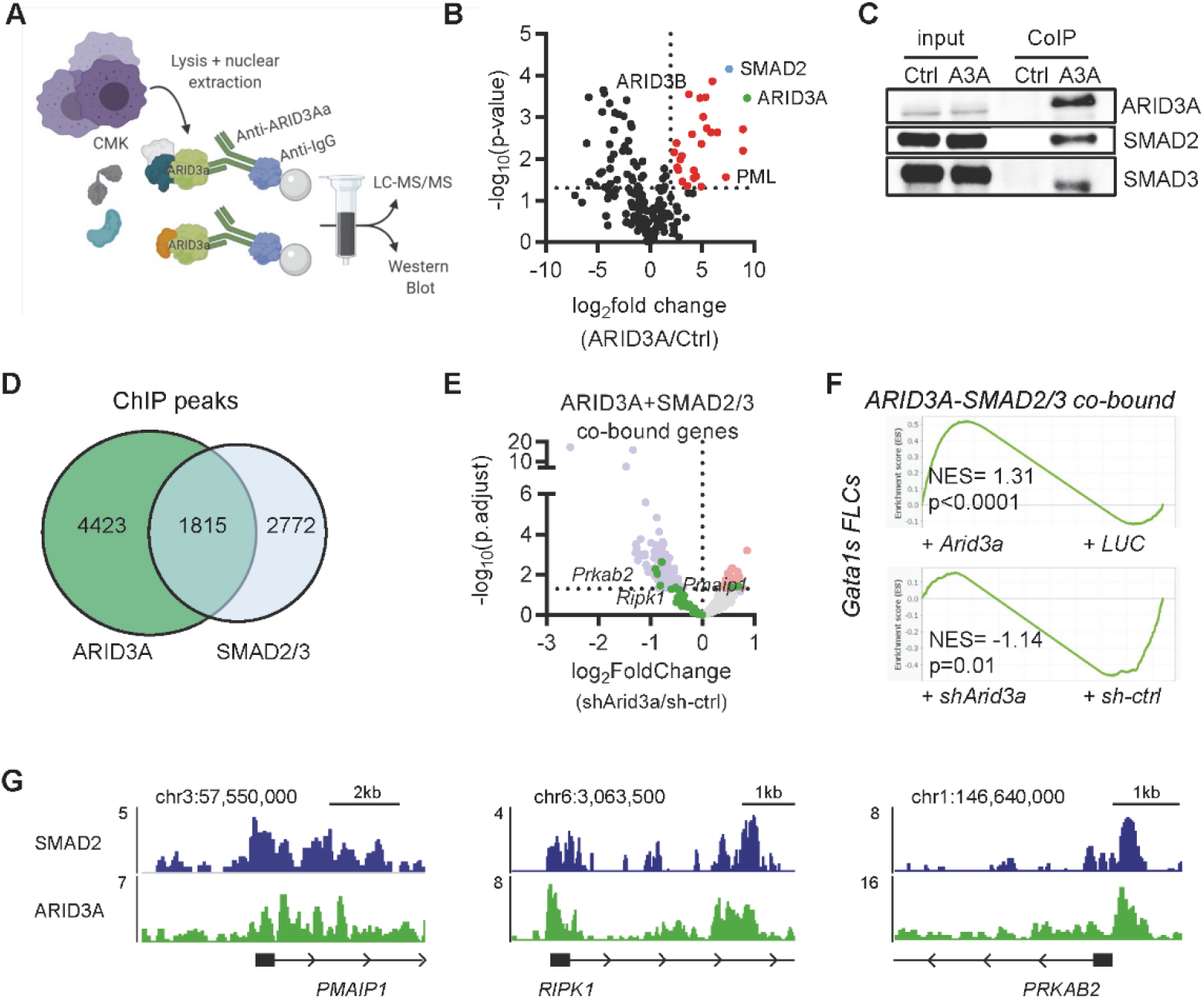
ARID3A interacts with SMAD2/3 and promotes TGFβ pathway activation. (A) Experimental design for isolating ARID3A-containing protein complexes from CMK cells. (B) Volcano plot showing enriched proteins in LC-MS/MS after ARID3A pulldown compared to the IgG control. Significantly-enriched proteins highlighted in red; ARID3A in green, SMAD2 in blue. (C) Western blot showing binding of SMAD2 and SMAD3 to co-immunoprecipitated ARID3A. (D) Venn diagram of ARID3A and SMAD2 and SMAD3 ChIP-seq peaks in K562 cells from the ENCODE datasets ^39^. (E) Volcano plot showing differential expression of genes bound by both ARID3A and GATA1 upon *Arid3a* knockdown in *Gata1s*-FLCs. Genes involved in apoptosis and cell cycle arrest are highlighted in green; significantly downregulated and upregulated genes in blue and red, respectively. (F) GSEA enrichment plots showing genes bound by ARID3A and SMAD2/3 (from D), and their response to *Arid3a* modulation in *Gata1s*-FLCs. (G) IGV snapshots of genes involved in apoptosis and cell cycle arrest that are co-bound and regulated by the ARID3A/SMAD2/3 protein complex (K562 cells, ENCODE datasets). Scale (y-axis) represents fold enrichment of normalized reads compared to the IgG control. Scale and chromosome location are shown (top).

In accordance with ARID3A’s previously described role in TGFβ signaling^50^, activation of TGFβ pathway genes correlated with Arid3a levels in *Gata1s*-FLCs (**Supplemental Figure 6A**). Moreover, we observed considerable co-occupancy of genomic loci by ARID3A and SMAD2/3 (29% of ARID3A occupied loci; 1815/6238) (**Figure 6D**), and expression of the co-bound genes – including several genes involved in cell cycle arrest and apoptosis such as *Pmaip1* (Noxa), *Ripk1* and *Prkab2*^51–53^ – correlated with *Arid3a* expression (**Figure 6E-G** and **Supplemental Figure 6B-C**). In line with these transcriptional changes, ectopic expression of *Arid3a* not only induced megakaryocytic differentiation (**Figure 3B**) but also led to rapid induction of apoptosis in *Gata1s*-FLCs overexpressing miR-125b, accompanied by a decrease in S-phase cells compared to the Luc control (**Supplemental Figure 6D-E**).

In summary, these results confirm the tumor suppressive role of ARID3A, positioning it as a mediator of TGFβ-induced cell cycle arrest and apoptosis in complex with SMAD2/3.

### Restoring ARID3A expression re-establishes normal differentiation of leukemic blasts

Gene expression data from AML patient samples show reduced *ARID3A* expression in TAM, ML-DS and non-DS AMKL compared to other AML subtypes and CD34^+^ HSPCs (**Figure 7A**). We therefore considered whether restoring *ARID3A* expression could be applied towards overcoming the leukemic phenotype and/or differentiation block in AMKL. Indeed, doxycycline-induced *ARID3A* expression in the ML-DS cell line CMK (**Supplemental Figure 7A**) led to growth arrest and a massive expansion of mature CD41^+^CD61^+^CD42b^+^ megakaryocytes, accompanied by induction of megakaryocytic genes as well as repression of the ML-DS expression signature and oncogenic programs (**Supplemental Figure 7B-E** and **Supplemental Table 8**). Corroborating its role as mediator of TGFβ signaling, we observed increased apoptosis and cell cycle arrest upon *ARID3A* expression (**Supplemental Figure 7F-G**), as well as reduced leukemic growth in other AMKL cell lines (**Supplemental Figure 7H**).

**Figure 7.**
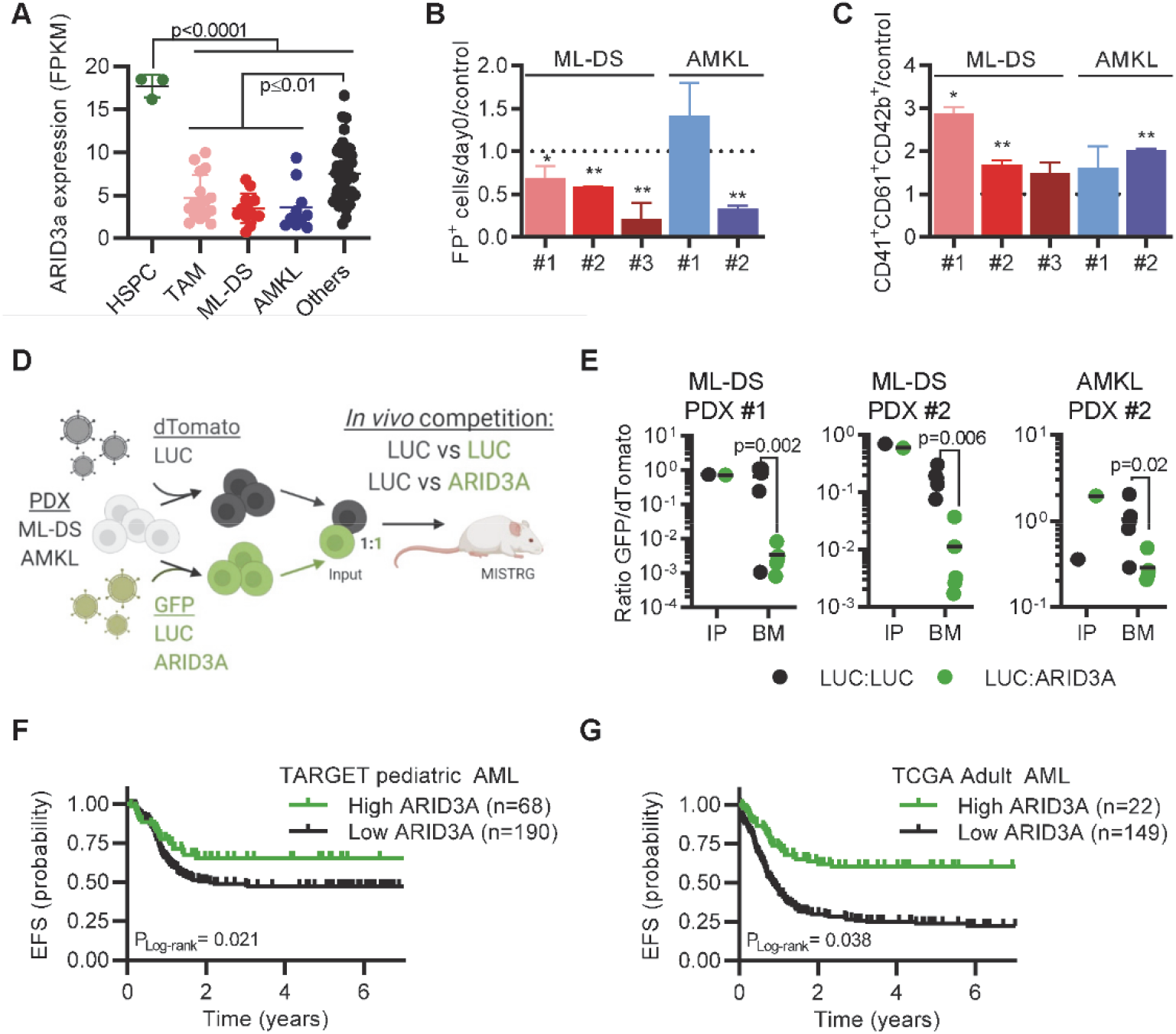
Restoring ARID3A expression re-establishes normal differentiation of leukemic blasts. (A) *ARID3A* expression (FPKM) in sorted pediatric AML blasts of different subtypes and CD34^+^ HSPCs. Others: inv(16), t(8;21), MLL-rearranged, t(10;11) and t(9;11) (unpaired t-test). (B-C) AMKL (n=2) and ML-DS (n=3) PDXs were transduced with doxycycline-inducible *ARID3A* or LUC cDNA vectors. (B) Bar graph showing the percentage of ARID3A^+^ cells after 12 days of induction with doxycycline, normalized to the LUC control. (C) Bar graph showing the percentage of ARID3A^+^ terminally differentiated megakaryocytic cells (CD41^+^CD61^+^CD42^+^) after 8 days of induction with doxycycline, normalized to the LUC control. *= p<0.05; ** = p<0.01 (n=3 per PDX, paired t-test). (D) Experimental design for evaluating whether ARID3A restores normal differentiation *in vivo*. Leukemic blasts were transduced with *ARID3A* (GFP^+^) or a LUC control (GFP^+^) and mixed 1:1 with LUC control-transduced blasts (dTomato^+^), before transplantation into sub-lethally irradiated recipient mice. (E) Ratio of GFP^+^ to dTomato^+^ cells in input cells (IP), and in the BM of mice sacrificed 4-5 weeks after transplantation (n=5, unpaired t-test). (F) Probability of event-free survival in 258 NCI-TARGET pediatric AML patients ^56^ with high (green; > 12.0 normalized reads; cutoff determined via maximally selected rank statistics) or low *ARID3A* expression (black; ≤ 12.0 normalized reads). (G) Probability of event-free survival in 171 TCGA adult AML patients ^57^ with high (green; > 12.3 normalized reads; cutoff determined via maximally selected rank statistics) or low *ARID3A* expression (black; ≤ 12.3 normalized reads). (A-E) All data are presented as mean ± SD.

We next restored *ARID3A* expression in ML-DS and non-DS AMKL patient samples that were expanded in mouse xenografts (see **Supplemental Table 2** for patient characteristics). Except for one non-DS AMKL sample harboring a *KMT2A* mutation, we observed a decrease in leukemic growth *in vitro* and induction of megakaryocytic differentiation upon doxycycline-induced *ARID3A* expression (**Figure 7B-C**). Lastly, we evaluated the effect of *ARID3A* re-expression on leukemic growth *in vivo* through a fluorescence-based competitive transplantation assay using two ML-DS and one AMKL patient-derived xenografts (PDX; **Figure 7D**). In all three cases, at the experimental endpoint, *ARID3A*-transduced leukemic blasts were significantly diminished in the bone marrow of the transplanted mice (**Figure 7E**), underlining the role of ARID3A as a miR-125b targeted novel tumor suppressor in ML-DS and AMKL. To assess the role of ARID3A in AML more globally, we analyzed the impact of *ARID3A* expression on the prognosis of AML across all subtypes and age groups. Kaplan-Meier analysis demonstrated that low *ARID3A* expression^54, 55^ was significantly associated with lower event-free survival and overall survival in a pediatric^56^ and adult^57^ cohort (**Figure 7G** and **Supplemental Figure 8A-D**), therefore identifying *ARID3A* expression as a prognostic factor. In line with a putative role as tumor suppressor in AML, ARID3A restoration exerted tumor suppressive effects in a panel of five myeloid leukemia cell lines *in vitro* and two PDXs *in vitro* and *in vivo,* representing AML subtypes with low ARID3A expression (**Supplemental Figure 8E-K**).

## Discussion

Here, we delineated the molecular basis of the differentiation blockade in AMKL. Using ML-DS – characterized by the interplay of trisomy 21 and *GATA1s* mutations – as a genetically simple model, we demonstrate that miR-125b increases proliferation and blocks terminal megakaryocytic differentiation in *Gata1s*-FLCs, thereby leading to leukemia *in vivo*, and that miR-125b-mediated downregulation of *Arid3a* underlies this phenotype. While the contribution of other miR-125b targets^20, 21, 58^ cannot be fully excluded, our comprehensive experimental data using complementary methods and different models strongly suggest *Arid3a* as the main target of miR-125b in this context. We further uncover novel functions for ARID3A in promoting megakaryocytic differentiation in concert with GATA1 and mediating TGFβ-induced cell cycle arrest and apoptosis in complex with SMAD2/3, indicating a tumor suppressive function for ARID3A in this context. Inversely, we propose that the dual hit of *ARID3A* repression and loss of full-length GATA1, and the resulting perturbation of erythropoiesis and megakaryopoiesis, is key to the pathogenesis of ML-DS. In this model, *Arid3a* downregulation blocks the otherwise unperturbed megakaryocytic differentiation of hyperproliferative *Gata1s* megakaryocytic progenitors, arresting them in an undetermined differentiation state and thereby causing leukemia. Reinstating *ARID3A* expression relieved megakaryocytic differentiation arrest in ML-DS and non-DS AMKL patient-derived xenografts, opening the way for new therapeutic concepts. As miR-125b is also upregulated in non-DS AMKL, which frequently presents with oncogene-mediated functional perturbation of GATA1 and acquired trisomy 21^59^, these findings are also more broadly applicable beyond the scope of TAM/ML-DS. This is underlined by our finding that *ARID3A* expression is a prognostic factor in AML across subtypes and age groups, which may be accounted to ARID3A’s function in controlling TGFβ signaling.

Our work on miR-125b and *Gata1s* represents a rare example of dissecting the specific interaction between a dysregulated miRNA and a known oncogene in the process of transformation. It was previously shown that *GATA1s* mutations perturb erythroid differentiation but leaves megakaryocytic differentiation unaffected^3, 4, 11, 12^. Accordingly, GATA1s occupancy at a number of erythroid specific genes is reduced but unaltered at megakaryocytic genes^3, 42, 60^. Since the erythroid and megakaryocytic lineages share a common progenitor, GATA1s has been speculated to bias differentiation away from the erythroid lineage and towards megakaryocytes, while failing to control their proliferation^21^. Here, we find that high levels of miR-125b – as seen in TAM and ML-DS – block this megakaryocytic route favored by *Gata1s* cells. Instead, the combination of *Gata1s* and miR-125b results in an accumulation of hyperproliferative megakaryocytic progenitors with erythroid features, which remain in an undetermined differentiation state and ultimately lead to leukemia *in vivo*. We note that the synergy between *Gata1s* and miR-125b alone in ML-DS was unexpected, as we previously showed that concerted action from all three members of the miR-99a~125b tricistron was required to enhance self-renewal and megakaryocytic differentiation in adult HSCs^20^. However, the current findings corroborate our previous study^21^ as well as others^61, 62^ describing miR-125b as an oncogenic miRNA in AML and ALL, and highlight the importance of developmental and cellular context in studying oncogenes and their partners – a phenomenon that has been described for other oncogenes in AMKL^59^.

Our results suggest an unexpected role for ARID3A in the regulation of megakaryopoiesis and megakaryocyte/erythroid cell fate decision, where high levels direct cells towards megakaryocytic and low levels towards erythroid fate. Our comprehensive molecular characterization of ARID3A further provides a mechanistic explanation for the synergy between miR-125b and GATA1s at the genetic level: ARID3A occupies megakaryocytic genes, inducing their transcriptional activation, and the majority of ARID3A regulated genes are also co-occupied by GATA1, suggesting that ARID3A acts as a GATA1 co-regulator during megakaryopoiesis. This co-regulation is sufficient to maintain megakaryocytic differentiation even in *GATA1s* mutated cells. Upon knockdown of *ARID3A,* this concerted transcriptional activation of megakaryocytic genes is lost, leading to a megakaryocytic differentiation arrest. Hence, together, *ARID3A* knockdown and *GATA1s* mutations block both functions of full-length *GATA1*, i.e. induction of erythroid and megakaryocytic differentiation. Since the respective abilities of GATA1 and ARID3A to control the proliferation of hematopoietic cells are similarly lost^63^, this combination results in drastic perturbation of normal hematopoiesis and development of megakaryoblastic leukemia with erythroid features *in vivo*, a hallmark of ML-DS^64, 65^. Using a proteomic approach, we found that the tumor suppressive functions of ARID3A are mediated in complex with SMAD2/3 – downstream effectors of the TGFβ signaling pathway, which induce apoptosis as well as cell cycle arrest in megakaryocytic progenitor cells^66, 67^. Thus, our current work adds the downregulation of *ARID3A* as another mechanism of miR-99a~125b-mediated control of TGFβ-signaling in hematopoietic cells^20^.

Overall, our works helps explain the synergy between trisomy 21 and *GATA1* mutations in leukemogenesis, through the functional dissection of a chromosome 21-encoded gene that is highly expressed in TAM/ML-DS. Although other factors on chromosome 21 – such as *CHAF1B^13^* and *RUNX1^15^* might be relevant for leukemogenesis, these findings can extend to non-DS AMKL with acquired trisomy 21 and functional GATA1 perturbation. In addition, we exemplify a general framework that can be used to interrogate oncogene-miRNA interactions in cancer and provide a basis for developing refined treatment approaches centered on *ARID3A*.

## Supporting information

Supplemental Methods, Supplemental Tables, Supplemental Figures and Supplemental References

## Acknowledgement

We thank D. Trono of EPFL, Lausanne, Switzerland, for kindly providing both pMD2.G (Addgene plasmid 12259) and psPAX2 (Addgene plasmid 12260). We thank Dr. A. Santos from the University of Halle-Wittenberg, Halle (Saale), Germany for support with flow cytometry. Illustrations (Fig 6A, Figure 7D and graphical abstract) were created with BioRender.com. K. Weigert is recipient of Mildred-Scheel doctoral research funding from the German Cancer Aid. This work was supported by funding to J.H.K from the German Research Foundation (DFG; KL2374/5-1) and the European Research Council (ERC) under the European Union’s Horizon 2020 research and innovation programme (grant agreement #714226). J.H.K. is a recipient of the St. Baldrick’s Robert J. Arceci Innovation Award. D.H is supported by the German Cancer Aid (#111743).

## Author contributions

O.A.V. performed experiments, analyzed the data and wrote the manuscript. K.W., M.N., S.E., C.B., M.F., H.I. and R.B. performed experiments, analyzed the data and revised the manuscript. M.L. performed experiments. E.R. and K.S. analyzed and interpreted the data and revised the manuscript. M.L.Y. supervised data analysis, interpreted the data and revised the manuscript. D.R. provided patient material and/or data and revised the manuscript. D.H. and J.H.K. designed the study, analyzed and interpreted the data, wrote the manuscript and academically drove the project.

