## Supplemental Methods, Supplemental Tables, Supplemental Figures and Supplemental References for "The megakaryocytic transcription factor ARID3A suppresses leukemia pathogenesis"

### Cell culture

Human cell lines were purchased from the German Collection of Microorganisms and Cell Cultures (DSMZ) and maintained according to the supplier's instructions. Adult CD34+ HSPCs were obtained from mobilized peripheral blood of anonymous healthy donors and enriched using anti-CD34 immunomagnetic microbeads. Media promoting megakaryocytic differentiation of HSPCs has been previously described[1]. Media promoting erythroid differentiation of HSPCs contained Stemspan SFEM II supplemented with 1% penicillin/streptomycin, 1% L-glutamine, 2µM dexamethasone, 1µM β-estradiol (Sigma Aldrich), 1U/ml EPO, 5ng/mL IL-3 and 12ng/mL SCF. Human PDXs were cultured in Stemspan SFEM II supplemented with 1% penicillin/streptomycin, 100ng/ml SCF, 100ng/ml FLT3-L, 20ng/ml IL-6, 50ng/ml TPO, 0.75µM StemRegenin1 and 35nM UM171. All cytokines were purchased from Peprotech. All cells were maintained at 37°C. Cells were transduced with concentrated viral particles in the presence of 2-5µg/ml polybrene. Primary human and murine HSPCs were transduced in Retronectin-coated plates according to manufacturer's instructions (Takara Bio). Murine Gata1s-FLCs were prepared and cultured as previously described [2] and cultured for at least 21 days. Day 0 was considered 96 hours post transduction or upon doxycycline addition (500ng/mL). Colony forming unit (CFU) assays, were performed as described previously[3,4], either in complete (HSC003 and HSC007, R&D Systems) or low (HSC006, R&D Systems; supplemented with 20ng mL Thpo) cytokine conditions. Micrographs were obtained with a Keyence BZ-9000 Microscope using the BZ-II viewer and processed with the BZ-II Analyzer software.

### Lentiviral production

Lentiviral particles were generated through the co-transfection of HEK293T cells with the corresponding expression constructs, pMD2.G and psPAX2 (Addgene #12259 and #12260), via the polyethyleneimine transfection method as previously described[3]. shRNAs were designed using the miR-N tool[5] applying the SENSOR design rules[6] and cloned into a SIN40C.SFFV.dTomato.miR30n backbone[7]. DNA sequences the different miRNAs[3] were cloned into a SIN40C.SFFV.miR30n backbone with different fluorescent proteins (dTomato, GFP and mTagBFP2, respectively). A non-silencing shRNA against Renilla luciferase in the miR-30n backbone was used as a control (sh-ctrl)[7]. Codon-optimized cDNAs were similarly cloned into the same SIN40C.SFFV backbone encoding GFP or dTomato as a fluorescent reporter. For inducible expression, a doxycycline-inducible SIN40C.TRE vector was used. All plasmids generated in this study have been deposited to Addgene and are listed in **Supplemental Table 1**.

### shRNA positive-selection screening

2 million Gata1s-FLCs were transduced with the miR-125b-mimicking shRNA library (MOI=0.3) to achieve sufficient representation (<500). Samples were harvested after 4 and 30 days in culture, and gDNA was extracted using the Blood Mini Kit (Qiagen). The shRNA amplicon was PCR-amplified using primers containing the p5 and p7 adaptor sequences, gel-purified and sequenced (single-end) with an Illumina HiSeq 2500. We used model-based analysis of genome-wide CRISPR/Cas9 knockout (MAGeCK)[20] to identify hits from the shRNA screen. Custom R scripts were used for demultiplexing double barcoded reads; guides with less than 20 reads in ≥75% of all samples were excluded. Raw read counts were passed to the mageck test command.

### Proteomic analysis

For total cell lysis and Western blotting previously described procedures were followed[4]. Blots were developed using Amersham™ ECL Prime Western Blotting Detection Reagent (Thermo Fisher Scientific). CoIP of endogenous ARID3A was performed on CMK cells using Anti-ARID3A antibody coupled to Novex™ DYNAL™ Dynabeads™ Protein G (Thermo Fisher Scientific). After cell lysis and affinity pulldown, proteins were eluted and subjected to proteolysis with trypsin (Promega) according to the filter-aided sample preparation (FASP) protocol[8]. Samples were analyzed by LC/MS/MS using a U3000 nano-HPLC system coupled to a Q-Exactive Plus mass spectrometer (Thermo Fisher Scientific). Raw data were processed using Proteome Discoverer 2.4 (Thermo Fisher Scientific). MS/MS data were searched against the Uniprot database (version Nov. 2019, tax. *Homo sapiens*, 73801 entries)[9] using Sequest-HT[10]. Label-free quantification of proteins was based on extracted peak areas of corresponding peptide precursor ions.

### 3'UTR binding assay

To evaluate the binding of miR-125b to the 3'UTR of the *ARID3A/Arid3a* mRNA, we cloned the 3'UTRs containing the miR-125b binding sites into a SIN40C.EFS.eGFP.pre lentiviral backbone downstream of the eGFP cassette. A 3'UTR where 3 nucleotides in the predicted miR-125b binding site were exchanged was used as mutated control. After lentiviral transduction, HEL cells were FACS-sorted and transduced with miR-125b or miR-control. Knockdown efficiency was defined as previously described[11].

### Flow cytometry and sorting

Flow cytometry was performed on a CytoFLEX Flow cytometer (Beckman Coulter) and the data analyzed in Kaluza 1.5 (Beckman Coulter). Antibodies are listed in **Supplemental Table S9**. Cell sorting was performed in a BD FACSAria™ II Flow Cytometer (BD Biosciences). Apoptosis was measured using the Annexin V Apoptosis Detection Kit II with APC-Annexin V (BD Biosciences) or PE-Cy7-Annexin V

(Thermo Fisher Scientific). Cell cycle analysis was performed with the BrdU Flow Kit (BD Biosciences) using Alexa Fluor 647 Anti-BrdU (BD Biosciences) or PE-Cy7 anti-BrdU (Biolegend). Both assays were performed according to manufacturers' instructions.

### Gene expression profiling

sgLuc- transduced FLCs were FACS-sorted three days after transduction; transduced CMK or Gata1s-FLCs were FACS-sorted after 2, 10 and 13 days of doxycycline induction. RNA was prepared using the Quick-RNA™ Miniprep Kit (Zymo Research). RNA-Sequencing was performed by Novogene UK. A minimum amount of 150ng RNA per sample was used as input material for the RNA sample preparations. Sequencing libraries were generated using NEBNext® UltraTM RNA Library Prep Kit for Illumina® (New England Biolabs) and sequenced on an Illumina NovaSeq using a PE150 system and paired-end reads were generated. Raw data (raw reads) of FASTQ format were firstly processed through fastp[12] and further processed as previously described[13]. Differential expression analysis was performed using DESeq2 R package[14]. The resulting P values were adjusted using the Benjamini and Hochberg's approach for controlling the False Discovery Rate (FDR)[15]. Genes with an adjusted P value <0.05 found by DESeq2 were assigned as differentially expressed. Calculation of functional enrichments was performed using gene set enrichment analysis (GSEA; v4.0)[16], including previously described and curated gene sets and ML-DS signatures[7]. Human gene symbols were mapped to murine gene symbols using orthologue annotations provided by Ensembl[17], considering only one-to-one orthologue relationships.

### Patient survival analysis

Event-free survival (EFS) was defined as time from diagnosis to the first event or last follow-up. Events were death from any cause, failure to achieve remission, relapse, and secondary malignancy. Failure to achieve remission was considered as an event on day 0. Overall survival was defined as the time between diagnosis and death from any cause or last follow-up. The Kaplan-Meier method was used to estimate survival rates[18]. Differences were compared using the 2-sided log-rank test[19], and standard errors were obtained using the Greenwood formula. The DESeq2 package was used to normalize and variance-stabilize RNA-sequencing read count data[14]. The pediatric AML data set further required batch correction, for which we used the sva package[20]. Normalized (and batch-corrected) expression of *ARID3A* was taken as a continuous variable in the survival model. For patient stratification, the optimal cutoff point was determined using maximally selected log-rank statistics as implemented in the maxstat R package (http://cran.r-project.org/web/packages/maxstat/index.html). The calculated cutoff for EFS was used for both overall survival and EFS analyses. We relied on R Version 3.6.1 (http://www.r-project.org/) for all of the above computations.

### Supplemental Tables

**Supplemental Table 1:** List of reagents and resources

**Supplemental Table 2**: Patient sample characteristics

**Supplemental Table 3**: Targets and sequences of the miR-125b-mimic shRNA library

**Supplemental Table 4**: Enrichment scores from the shRNA-based positive selection screen

**Supplemental Table 5**: GSEA results of global gene expression profiling after overexpression of miR-125b in *Gata1s*-FLCs

**Supplemental Table 6**: List of shRNA targeting *ARID3A*/*Arid3a*

**Supplemental Table 7**: GSEA results of global gene expression profiling after modulation of *Arid3a* in *Gata1s*-FLCs

**Supplemental Table 8**: Pairwise analysis of LC-MS/MS

**Supplemental Table 9**: GSEA results of global gene expression profiling after overexpression of *ARID3A* in CMK

**Supplemental Table 2**: Patient sample characteristics

| | gender | age at diagnosis (years) | WBC (x109/L) | hemoglobin (g/dl) | BM blasts (%) | CNS | SCT | molecular genetics | Cytogenetics (karyotype) | response | relapse |
|---|---|---|---|---|---|---|---|---|---|---|---|
| ML-DS PDX#1 | m | 2 1/6 | 32500 | 7.9 | 70.5 | no | yes | GATA1 mutation | 47,XY,t(3;13)(q?26;q?13~14) del(13)(q?14q22),+21c[cp14]/ 47,sl,del?(15)(q?)[cp2]/46,XY[1] | NR, CCR:after SCT | no |
| ML-DS PDX#2 | m | 1 1/3 | 4800 | 12 | 10.5 | no | no | GATA1 mutation | k.A. | CCR | no |
| ML-DS PDX#3 | f | 1 1/12 | 168000 | 7.1 | 16 | no | no | NRAS mutation | k.A. | CCR | no |
| AMKL PDX#1 | m | 1 | 40000 | 9.9 | 64 | no | yes | KMT2A mutation | 46,XY[15].nuc ish 3q26(EVI1x2)[100/100], 8q22(RUNX1T1x2),21q22(RUNX1x2)[98/100], 11 q23(MLLx2)[99/100], 16q22(CBFBx2)[100/100] 17q21.1(RARAx2)[100/100] | CCR | yes |
| AMKL PDX#2 | m | 2/12 | 33800 | 9.5 | 34 | yes | no | n.d. | 46,XY,t(1;8;22)(p13;q22;q13)[14]/46,XY[1] | NR | no |

**Supplemental Table 4**: Enrichment scores from the shRNA-based positive selection screen

| Gene ID | $log_2$fold change | p.value | FDR | Gene ID | $log_2$fold change | p.value | FDR |
|---|---|---|---|---|---|---|---|
| Podxl | 18,211 | 0,003266 | 0,692389 | Tdg | -0,61991 | 0,51773 | 0,978994 |
| Reep3 | 13,431 | 0,0091298 | 0,74172 | Fam129b | -0,48297 | 0,53076 | 0,978994 |
| Lpp | 13,446 | 0,010496 | 0,74172 | Slc38a9 | -0,28405 | 0,53416 | 0,978994 |
| E2f3 | 19,416 | 0,017347 | 0,744949 | Acvr2a | -0,02331 | 0,53563 | 0,978994 |
| Fzd5 | 18,417 | 0,021825 | 0,744949 | Mknk2 | -0,41859 | 0,54057 | 0,978994 |
| Unc13b | 11,016 | 0,022882 | 0,744949 | Zswim4 | -0,28989 | 0,54357 | 0,978994 |
| Phc2 | 19,856 | 0,028716 | 0,744949 | Otub2 | -0,46519 | 0,55007 | 0,978994 |
| Rnf144b | 12,311 | 0,030172 | 0,744949 | Arid3b | -0,70642 | 0,55106 | 0,978994 |
| Arid3a | 10,512 | 0,03808 | 0,744949 | Ren | -0,23496 | 0,5538 | 0,978994 |
| Map3k3 | 10,512 | 0,040254 | 0,744949 | Gas2l1 | -0,4527 | 0,55618 | 0,978994 |
| Mef2c | 0,79338 | 0,043784 | 0,744949 | Fgfr1 | 0,13884 | 0,55941 | 0,978994 |
| Bmf | 0,65678 | 0,046188 | 0,744949 | Plb1 | 0,21637 | 0,5625 | 0,978994 |
| Cdr2l | 14,542 | 0,047075 | 0,744949 | Tmem | 0,17315 | 0,58004 | 0,978994 |
| Rbm38 | 0,30738 | 0,053767 | 0,744949 | Accn2 | -0,35256 | 0,58119 | 0,978994 |
| Slc8a2 | 0,98054 | 0,057307 | 0,744949 | Dapk1 | -0,37717 | 0,58309 | 0,978994 |
| Ptpro | -0,27504 | 0,060009 | 0,744949 | Klf13 | -0,22224 | 0,59137 | 0,978994 |
| Mgat4a | 0,79538 | 0,061575 | 0,744949 | Ganc | -0,64927 | 0,59447 | 0,978994 |
| Trim71 | 14,452 | 0,06325 | 0,744949 | Pik3ip1 | -0,09572 | 0,59794 | 0,978994 |
| Acer3 | 0,32946 | 0,073492 | 0,800053 | Neu1 | -0,15549 | 0,60369 | 0,978994 |
| AB124611 | 10,131 | 0,075477 | 0,800053 | Cpeb3 | 0,090764 | 0,60671 | 0,978994 |
| Setd7 | 0,3349 | 0,084332 | 0,812389 | Tet2 | -0,06904 | 0,60934 | 0,978994 |
| Atp13a3 | 0,30916 | 0,086028 | 0,812389 | Rit1 | 0,012464 | 0,6291 | 0,978994 |
| 1700025G04Rik | 12,827 | 0,088281 | 0,812389 | Tnfrsf1b | -0,60858 | 0,63755 | 0,978994 |
| Abl2 | 0,64403 | 0,093686 | 0,812389 | Tmem26 | -0,34392 | 0,63984 | 0,978994 |
| Gcnt1 | 0,82238 | 0,095801 | 0,812389 | Bach1 | -0,21152 | 0,65013 | 0,978994 |
| Il13 | 0,60133 | 0,11553 | 0,902635 | Arhgef1 | -0,61972 | 0,65849 | 0,978994 |
| Rhot2 | 11,383 | 0,11607 | 0,902635 | Sh2b3 | -0,72887 | 0,66269 | 0,978994 |
| Ctdsp2 | 0,20944 | 0,11922 | 0,902635 | Trib2 | -0,7399 | 0,66931 | 0,978994 |
| Abhd3 | 0,70079 | 0,13018 | 0,937048 | Slco2b1 | -0,53728 | 0,67162 | 0,978994 |
| Foxn3 | 0,38527 | 0,13524 | 0,937048 | Celsr2 | -0,49381 | 0,67594 | 0,978994 |
| Zfp697 | 0,60179 | 0,13974 | 0,937048 | Bcat1 | -0,69375 | 0,6822 | 0,978994 |
| Tnfaip8l3 | 0,32358 | 0,15138 | 0,937048 | Npl | -0,79773 | 0,68688 | 0,978994 |
| Rufy3 | 0,62091 | 0,15338 | 0,937048 | Vps4b | -0,13009 | 0,68758 | 0,978994 |
| pGL2 | 0,78786 | 0,15853 | 0,937048 | Tmod2 | -0,92995 | 0,68873 | 0,978994 |
| Ankrd29 | 0,64589 | 0,15892 | 0,937048 | Coro1c | -0,78896 | 0,69119 | 0,978994 |
| Apaf1 | 0,22716 | 0,16527 | 0,937048 | Adam9 | -0,11693 | 0,6954 | 0,978994 |
| Itga7 | -0,49822 | 0,16535 | 0,937048 | Mapk14 | -0,6026 | 0,69766 | 0,978994 |
| Dusp3 | 0,80461 | 0,16796 | 0,937048 | Mknk1 | -0,26853 | 0,69821 | 0,978994 |
| Mtus1 | 0,68293 | 0,18136 | 0,978994 | Acvr2b | -0,86558 | 0,70004 | 0,978994 |
| Dram2 | 0,49792 | 0,19822 | 0,978994 | Trps1 | -0,74998 | 0,70081 | 0,978994 |
| Prdm1 | -0,43376 | 0,20856 | 0,978994 | Wipf2 | 0,11111 | 0,70418 | 0,978994 |

| | | | | | | | |
|---|---|---|---|---|---|---|---|
| Mycl1 | 0,43842 | 0,22258 | 0,978994 | Traf6 | -0,38773 | 0,70536 | 0,978994 |
| Syn2 | 0,61753 | 0,22503 | 0,978994 | Irf4 | -0,29991 | 0,70783 | 0,978994 |
| Tmem87b | 0,55059 | 0,23518 | 0,978994 | Abhd6 | -0,12295 | 0,71021 | 0,978994 |
| Tbc1d14 | 0,2559 | 0,23784 | 0,978994 | Snx18 | -0,88071 | 0,71383 | 0,978994 |
| Lactb | 0,40593 | 0,24138 | 0,978994 | Mcl1 | -0,92353 | 0,7223 | 0,978994 |
| Actr10 | 0,6169 | 0,24384 | 0,978994 | Clic4 | -10,906 | 0,72821 | 0,978994 |
| Socs4 | 0,49086 | 0,2459 | 0,978994 | Sgpl1 | -0,82952 | 0,73709 | 0,978994 |
| Rap1gap2 | -0,11252 | 0,25879 | 0,978994 | Baz2a | -0,72397 | 0,7395 | 0,978994 |
| Tlr8 | 0,45459 | 0,26193 | 0,978994 | St8sia4 | -0,89706 | 0,74422 | 0,978994 |
| Synj2bp | -0,7179 | 0,26485 | 0,978994 | 4930506M07Rik | -0,19725 | 0,74764 | 0,978994 |
| Myo1e | 0,33647 | 0,27106 | 0,978994 | Cyth1 | -0,75454 | 0,75325 | 0,978994 |
| Zfp191 | -0,042509 | 0,27414 | 0,978994 | Stard13 | -0,66165 | 0,75402 | 0,978994 |
| Rbm20 | 0,42573 | 0,27709 | 0,978994 | Apc | -0,73266 | 0,75865 | 0,978994 |
| Ap4e1 | 0,4165 | 0,27825 | 0,978994 | Csnk2a1 | -0,47077 | 0,76303 | 0,978994 |
| Tgfbr1 | 0,34813 | 0,28528 | 0,978994 | Slc26a6 | -1,281 | 0,769 | 0,978994 |
| Srd5a3 | 0,44488 | 0,28654 | 0,978994 | Rhoq | -1,211 | 0,77629 | 0,978994 |
| Slc25a24 | 0,34794 | 0,29272 | 0,978994 | Tmem86a | -0,86469 | 0,788 | 0,978994 |
| Ube2g1 | 0,368 | 0,29547 | 0,978994 | Cpeb4 | -0,28298 | 0,79291 | 0,978994 |
| Map3k1 | 0,080211 | 0,29665 | 0,978994 | Rab8b | -0,39113 | 0,7985 | 0,978994 |
| Nrp2 | 0,36459 | 0,29716 | 0,978994 | Vps36 | -0,31183 | 0,80079 | 0,978994 |
| Pou2f2 | 0,29057 | 0,29933 | 0,978994 | Glipr1 | -0,76585 | 0,80981 | 0,978994 |
| Lipa | 0,25547 | 0,30457 | 0,978994 | Etv6 | -0,29548 | 0,81128 | 0,978994 |
| Dnajb5 | -0,10121 | 0,31046 | 0,978994 | Lin28b | -0,7023 | 0,81554 | 0,978994 |
| Sash1 | -0,34621 | 0,31667 | 0,978994 | Limd1 | -0,77919 | 0,82361 | 0,978994 |
| Lbh | -0,26441 | 0,32523 | 0,978994 | Slc38a1 | -0,27681 | 0,8255 | 0,978994 |
| Trp53inp1 | -0,21259 | 0,33128 | 0,978994 | Smarcd1 | -15,139 | 0,82567 | 0,978994 |
| Asb13 | -0,078531 | 0,33426 | 0,978994 | Slc16a6 | -0,94919 | 0,82993 | 0,978994 |
| Mllt4 | -0,24686 | 0,33428 | 0,978994 | Myo7a | -0,63746 | 0,84056 | 0,978994 |
| Msr1 | -0,46842 | 0,3402 | 0,978994 | Il6ra | -0,68388 | 0,8427 | 0,978994 |
| Pald1 | 0,25211 | 0,34725 | 0,978994 | Klf3 | -0,19003 | 0,84834 | 0,978994 |
| Frmd4b | 0,31737 | 0,3548 | 0,978994 | Borcs6 | -11,788 | 0,84872 | 0,978994 |
| Zfp710 | -0,55893 | 0,35822 | 0,978994 | Qk | -0,71527 | 0,85115 | 0,978994 |
| Rtcd1 | 0,20179 | 0,35836 | 0,978994 | St6gal1 | -0,31826 | 0,85178 | 0,978994 |
| Smad2 | -0,26605 | 0,36126 | 0,978994 | Rab22a | -0,79094 | 0,85311 | 0,978994 |
| Mob3b | 0,25667 | 0,36238 | 0,978994 | Tmem135 | -11,746 | 0,85781 | 0,978994 |
| Map2k7 | -0,50981 | 0,36447 | 0,978994 | Fam65b | -0,56516 | 0,86327 | 0,978994 |
| Ccr7 | 0,13359 | 0,36734 | 0,978994 | Cgn | -13,818 | 0,86705 | 0,978994 |
| Mllt10 | 0,014767 | 0,37345 | 0,978994 | Fmn1 | -0,2668 | 0,87236 | 0,978994 |
| Rora | -0,27941 | 0,37913 | 0,978994 | Olfml2a | -0,60646 | 0,87556 | 0,978994 |
| Ncor2 | 0,1767 | 0,38154 | 0,978994 | Gpc4 | -0,7572 | 0,87726 | 0,978994 |
| Lrrk2 | 0,094665 | 0,38844 | 0,978994 | Mapk12 | -11,806 | 0,87834 | 0,978994 |
| Zswim6 | -0,14512 | 0,3916 | 0,978994 | Arrdc4 | -0,76514 | 0,88483 | 0,978994 |
| Psmb8 | -0,31479 | 0,39451 | 0,978994 | Ppp2ca | -0,68708 | 0,88644 | 0,978994 |
| Cdk19 | -0,58488 | 0,39729 | 0,978994 | Trim7 | -13,032 | 0,88796 | 0,978994 |
| Slc16a10 | -0,48462 | 0,4005 | 0,978994 | Anapc16 | -14,396 | 0,89237 | 0,978994 |

| | | | | | | | |
|---|---|---|---|---|---|---|---|
| Nt5dc1 | 0,03504 | 0,40385 | 0,978994 | Slc46a3 | -0,95053 | 0,89565 | 0,978994 |
| Bmpr2 | 0,23971 | 0,42228 | 0,978994 | Pi4k2a | -1,07 | 0,89857 | 0,978994 |
| Ank | -0,024537 | 0,42976 | 0,978994 | Rassf3 | -13,508 | 0,90049 | 0,978994 |
| Bcl2l12 | 0,21234 | 0,43416 | 0,978994 | Klhl24 | -0,49806 | 0,90619 | 0,980169 |
| Abtb1 | -0,17867 | 0,44033 | 0,978994 | Sept11 | -14,237 | 0,91479 | 0,984444 |
| Maf | 0,25079 | 0,44511 | 0,978994 | Dus1l | -1,315 | 0,92243 | 0,984962 |
| Sertad3 | -0,32769 | 0,4597 | 0,978994 | Baiap2 | -10,806 | 0,92456 | 0,984962 |
| Atxn1l | -0,36812 | 0,46569 | 0,978994 | Plekhm3 | -0,52546 | 0,93347 | 0,986029 |
| pGL3-Luc | -0,059947 | 0,48111 | 0,978994 | Daam2 | -17,783 | 0,94205 | 0,986029 |
| Dusp7 | -0,13643 | 0,48166 | 0,978994 | Dazap2 | -0,80727 | 0,9421 | 0,986029 |
| Mfsd7c | -0,31089 | 0,48186 | 0,978994 | Ttc7 | -13,897 | 0,94417 | 0,986029 |
| Tab2 | -0,44637 | 0,48491 | 0,978994 | Homez | -10,914 | 0,95449 | 0,987826 |
| 1700017B05Rik | -0,1604 | 0,48626 | 0,978994 | Ppm1f | -1,284 | 0,95676 | 0,987826 |
| Pctp | -0,22003 | 0,48808 | 0,978994 | D17H6S53E | -13,874 | 0,96014 | 0,987826 |
| Pik3c2b | -0,16198 | 0,49288 | 0,978994 | Naif1 | -21,675 | 0,96739 | 0,987826 |
| E2f2 | 0,28522 | 0,49397 | 0,978994 | Ipmk | -0,91672 | 0,97375 | 0,987826 |
| Pdpk1 | -0,017022 | 0,49701 | 0,978994 | Snx30 | -22,969 | 0,97385 | 0,987826 |
| St5 | 0,20124 | 0,50954 | 0,978994 | Rxra | -12,655 | 0,98608 | 0,994835 |
| Srgap2 | -0,39678 | 0,5125 | 0,978994 | Kif1b | -16,994 | 0,99014 | 0,994835 |
| Cecr6 | 0,097405 | 0,51548 | 0,978994 | Ahrr | -19,762 | 0,99713 | 0,997133 |

**Supplemental Table 5**: GSEA results of global gene expression profiling after overexpression of miR-125b in *Gata1s*-FLCs

**Hallmark gene sets** [22]

| # | Gene set | NES | NOM p | FDR q |
|---|---|---|---|---|
| 1 | HALLMARK_OXIDATIVE_PHOSPHORYLATION | 2,292 | 0,000 | 0,000 |
| 2 | HALLMARK_MYC_TARGETS_V1 | 2,279 | 0,000 | 0,000 |
| 3 | HALLMARK_MYC_TARGETS_V2 | 1,916 | 0,000 | 0,000 |
| 4 | HALLMARK_DNA_REPAIR | 1,679 | 0,000 | 0,004 |
| 5 | HALLMARK_E2F_TARGETS | 1,591 | 0,000 | 0,009 |
| 6 | HALLMARK_FATTY_ACID_METABOLISM | 1,392 | 0,009 | 0,042 |
| 7 | HALLMARK_REACTIVE_OXYGEN_SPECIES_PATHWAY | 1,347 | 0,040 | 0,058 |
| 8 | HALLMARK_UV_RESPONSE_UP | 1,324 | 0,000 | 0,068 |
| 9 | HALLMARK_ADIPOGENESIS | 1,303 | 0,006 | 0,072 |
| 10 | HALLMARK_MTORC1_SIGNALING | 1,251 | 0,025 | 0,104 |
| 11 | HALLMARK_KRAS_SIGNALING_DN | -1,450 | 0,001 | 0,148 |
| 12 | HALLMARK_MITOTIC_SPINDLE | -1,317 | 0,011 | 0,394 |
| 13 | HALLMARK_HEDGEHOG_SIGNALING | -1,285 | 0,091 | 0,382 |
| 14 | HALLMARK_HEME_METABOLISM | -1,231 | 0,054 | 0,514 |
| 15 | HALLMARK_TNFA_SIGNALING_VIA_NFKB | -1,222 | 0,050 | 0,451 |

**StemCell and matured lineages genesets** [7]

| # | Gene set | NES | NOM p | FDR |
|---|---|---|---|---|
| 1 | WONG_EMBRYONIC_STEM_CELL_CORE | 2,050 | 0,000 | 0,000 |
| 2 | EZH2-KO IN ETP_UP | 1,749 | 0,000 | 0,013 |
| 3 | EZH2-KO IN ETP_IN_VIVO_UP | 1,673 | 0,000 | 0,021 |
| 4 | BHATTACHARYA_EMBRYONIC_STEM_CELL | 1,633 | 0,000 | 0,027 |
| 5 | ES_MYC_MODULE | 1,585 | 0,000 | 0,035 |
| 6 | LU_EZH2_TARGETS_UP | 1,522 | 0,000 | 0,050 |
| 7 | GEORGOPOLOUS_MYELOID DIFFERENTIATION | 1,424 | 0,000 | 0,101 |
| 8 | LAURENTI_CMP | 1,349 | 0,000 | 0,165 |
| 9 | LAURENTI_GMP | 1,320 | 0,000 | 0,185 |
| 10 | KUSTIKOVA TOP200_DOWN_EVI1 | 1,303 | 0,010 | 0,187 |
| 11 | GEORGOPOLOUS_GMP VS ALL OTHER +CD33 | 1,300 | 0,024 | 0,174 |
| 12 | ROSS_AML_WITH_MLL_FUSIONS | 1,276 | 0,062 | 0,191 |
| 13 | REGEV_G1-S_CORE SET | 1,273 | 0,090 | 0,180 |
| 14 | EBERT_HUMAN_MYELOID | 1,263 | 0,012 | 0,182 |
| 15 | KLUSMANN_ARRAYSTAR_MONO | 1,259 | 0,058 | 0,175 |
| 16 | HIDALGO_EZH1-KO IN HSC_UP | 1,250 | 0,034 | 0,176 |
| 17 | LAURENTI_MEP | 1,235 | 0,000 | 0,188 |
| 18 | KLUSMANN_NCODE_CD34 | 1,222 | 0,040 | 0,198 |
| 19 | KLUSMANN_ARRAYSTAR_ERYTHROID | 1,080 | 0,211 | 0,405 |
| 20 | GOODELL_DNMT3 TARGETS | -1,594 | 0,000 | 0,062 |
| 21 | IWAMA_EZH2-KO IN HSC_FL_UP | -1,476 | 0,000 | 0,186 |

| | | | | |
|---|---|---|---|---|
| 22 | EPPERT_LSC-R_EXTENDED | -1,445 | 0,009 | 0,191 |
| 23 | EBERT_HUMAN_LYMPHOID | -1,421 | 0,017 | 0,194 |
| 24 | EBERT_HUMAN_TCELLS | -1,418 | 0,014 | 0,161 |
| 25 | KLUSMANN_NCODE_GRANULOCYTES | -1,330 | 0,015 | 0,379 |
| 26 | EPPERT_LSC-R | -1,323 | 0,083 | 0,347 |
| 27 | KUSTIKOVA TOP250_UP_EVI1 | -1,289 | 0,010 | 0,438 |
| 28 | KLUSMANN_NCODE_MEGAKARYOCYTIC | -1,267 | 0,042 | 0,492 |
| 29 | ROSSI_T-CELLS | -1,261 | 0,047 | 0,473 |
| 30 | FISCHER_DOWN IN RC | -1,246 | 0,080 | 0,499 |
| 31 | KONDO_EZH2_TARGETS | -1,240 | 0,045 | 0,484 |
| 32 | LAURENTI_MLP | -1,235 | 0,046 | 0,467 |
| 33 | LU_EZH2_TARGETS_DN | -1,215 | 0,032 | 0,523 |
| 34 | LAURENTI_HSC1 AND HSC2 | -1,08 | 0,273 | 0,614 |

**ML-DS signature genesets** [7]

| # | Gene set | NES | NOM p | FDR q |
|---|---|---|---|---|
| 1 | ML-DS TOP600 UP.GRP | 1,899 | 0,000 | 0,000 |
| 2 | ML-DS TOP500 UP.GRP | 1,847 | 0,000 | 0,000 |
| 3 | ML-DS TOP400 UP.GRP | 1,842 | 0,000 | 0,000 |
| 4 | ML-DS TOP250 UP.GRP | 1,772 | 0,000 | 0,000 |
| 5 | ML-DS TOP300_UP GENE.GRP | 1,760 | 0,000 | 0,000 |
| 6 | ML-DS TOP100_UP GENE.GRP | 1,641 | 0,000 | 0,000 |
| 7 | ML-DS TOP100_DOWN GENE.GRP | 1,043 | 0,347 | 0,278 |
| 8 | ML-DS TOP300_DOWN GENE.GRP | -1,246 | 0,049 | 0,227 |

**Supplemental Table 6**: List of shRNA targeting *ARID3A*/*Arid3a*

| **Mouse** | **Sequence** |
|---|---|
| shArid3a #1 | CTAGGAAGAATCTATCTGTATATAGTGAAGCCACAGATGTATATACAGATAGATTCTTCCTAA |
| shArid3a #2 | AAAGAATCTATCTGTATATCTATAGTGAAGCCACAGATGTATAGATATACAGATAGATTCTTC |
| shArid3a #3 | ACCCAAGATCAAGAAAGAGGAATAGTGAAGCCACAGATGTATTCCTCTTTCTTGATCTTGGGC |
| shArid3a #4 | AAGGACACAGAATGTTCTAGAATAGTGAAGCCACAGATGTATTCTAGAACATTCTGTGTCCTG |

| **Human** | **Sequence** |
|---|---|
| shARID3A #1 | CAAGGATATCTATATATCTATATAGTGAAGCCACAGATGTATATAGATATATAGATATCCTTT |
| shARID3A #2 | AGCCGATCCTGTTTACCTCATATAGTGAAGCCACAGATGTATATGAGGTAAACAGGATCGGCC |
| shARID3A #3 | ATGGATGACTTGTTCAGCTTCATAGTGAAGCCACAGATGTATGAAGCTGAACAAGTCATCCAG |

| **Non-targeting** | **Sequence** |
|---|---|
| shLUC | CCAGGATTACAAGATTCAAAGTTAGTGAAGCCACAGATGTAACTTTGAATCTTGTAATCCTGA |

**Supplemental Table 8**: Pairwise analysis of LC-MS/MS

| Protein | log2FC (ARID3A/Control) | $log_2$pvalue |
|---|---|---|
| SMAD2 | 7,62 | 4,159926 |
| P4HA1 | 6,01 | 3,867153 |
| ARID3B | 3,73 | 3,555343 |
| TRIM21 | 5,36 | 3,488007 |
| RBM15 | 4,82 | 3,466823 |
| ARID3A | 9,33 | 3,465567 |
| SERPINH1 | 5,12 | 3,011139 |
| SRRM2 | 5,48 | 2,736189 |
| DACH1 | 8,93 | 2,718777 |
| P4HB | 6,49 | 2,636633 |
| PCM1 | 5,85 | 2,634229 |
| DHX15 | 4,05 | 2,5961 |
| RNPS1 | 2,55 | 2,382591 |
| MIB1 | 4,96 | 2,362444 |
| COL1A1 | 8,94 | 2,196879 |
| HSPA8 | 2,33 | 2,160015 |
| RPL3 | 3,06 | 2,074202 |
| NUMA1 | 2,93 | 1,990553 |
| SSR4 | 2,6 | 1,796137 |
| HADHA | 2,75 | 1,730748 |
| CCT7 | 4,21 | 1,723534 |
| XRCC5 | 4,44 | 1,579291 |
| PML | 7,3 | 1,567755 |
| LRRC59 | 4,21 | 1,539919 |
| KPNB1 | 3,11 | 1,455751 |
| RAN | 3,59 | 1,349624 |
| DLST | 4,94 | 1,340969 |
| PKM | 3,7 | 1,179455 |
| XRCC6 | 4,12 | 1,17568 |
| VDAC1 | 2,56 | 0,860005 |
| SSBP1 | 2,67 | 0,742064 |
| ENO1 | 3,45 | 0,592388 |
| A2ML1 | 2,64 | 0,489387 |
| EEF2 | 2,29 | 0,390247 |
| VCP | 2,28 | 0,309687 |
| GAPDH | 2,36 | 0,217776 |
| GSTP1 | 2,78 | 0,124319 |

**Supplemental Table 9**: GSEA results of global gene expression profiling after overexpression of *ARID3A* in CMK

**Hallmark gene sets**[22]

| # | Gene set | NES | NOM p | FDR q |
|---|---|---|---|---|
| 1 | HALLMARK_TGF_BETA_SIGNALING | 1,478 | 0,011 | 0,105 |
| 2 | HALLMARK_MYOGENESIS | 1,416 | 0,003 | 0,146 |
| 3 | HALLMARK_TNFA_SIGNALING_VIA_NFKB | 1,357 | 0,011 | 0,232 |
| 4 | HALLMARK_P53_PATHWAY | 1,330 | 0,020 | 0,249 |
| 5 | HALLMARK_KRAS_SIGNALING_DN | 1,280 | 0,035 | 0,376 |
| 6 | HALLMARK_E2F_TARGETS | -1,636 | 0,000 | 0,017 |
| 7 | HALLMARK_MYC_TARGETS_V1 | -1,625 | 0,000 | 0,010 |
| 8 | HALLMARK_UNFOLDED_PROTEIN_RESPONSE | -1,301 | 0,023 | 0,181 |
| 9 | HALLMARK_MYC_TARGETS_V2 | -1,140 | 0,188 | 0,494 |
| 10 | HALLMARK_G2M_CHECKPOINT | -1,074 | 0,207 | 0,353 |

**StemCell and matured lineages genesets**[7]

| # | Gene set | NES | NOM p | FDR |
|---|---|---|---|---|
| 1 | KLUSMANN_ARRAYSTAR_MEGA | 1,584 | 0,000 | 0,029 |
| 2 | EZH2-KO_IN ETP_H3K27ME3_DOWN | 1,580 | 0,000 | 0,015 |
| 3 | LAURENTI_MLP | 1,496 | 0,000 | 0,071 |
| 4 | KLUSMANN_NCODE_NKC | 1,471 | 0,006 | 0,089 |
| 5 | KLUSMANN_NCODE_MEGAKARYOCYTIC | 1,356 | 0,005 | 0,427 |
| 6 | IWAMA_EZH2-KO IN HSC_BM_UP | 1,324 | 0,024 | 0,519 |
| 7 | KUSTIKOVA TOP400_UP_EVI1 | 1,317 | 0,006 | 0,489 |
| 8 | KLUSMANN_ARRAYSTAR_HSPC_CB VS ALL_TOP200 | 1,308 | 0,037 | 0,473 |
| 9 | GOODELL_NK | 1,287 | 0,095 | 0,528 |
| 10 | GOODELL_NAIVE | 1,283 | 0,115 | 0,500 |
| 11 | KLUSMANN_NCODE_GRANULOCYTES | 1,276 | 0,037 | 0,489 |
| 12 | KLUSMANN_ARRAYSTAR_GRANULOCYTES | 1,264 | 0,036 | 0,509 |
| 13 | LAURENTI_HSC1 VS HSC2 | 1,004 | 0,482 | 0,719 |
| 14 | REGEV_G1-S_CORE SET | -1,748 | 0,000 | 0,005 |
| 15 | LAURENTI_MEP | -1,744 | 0,000 | 0,003 |
| 16 | ROSS_AML_OF_FAB_M7_TYPE | -1,700 | 0,000 | 0,004 |
| 17 | KAMMINGA_EZH2_TARGETS | -1,519 | 0,000 | 0,052 |
| 18 | JAATINEN_HEMATOPOIETIC_STEM_CELL_UP | -1,488 | 0,000 | 0,061 |
| 19 | EBERT_HUMAN_CD34 | -1,463 | 0,000 | 0,066 |
| 20 | WONG_EMBRYONIC_STEM_CELL_CORE | -1,427 | 0,000 | 0,079 |
| 21 | EBERT_HUMAN HSC | -1,357 | 0,012 | 0,124 |
| 22 | KLUSMANN_ARRAYSTAR_MONO | -1,343 | 0,031 | 0,121 |
| 23 | KUSTIKOVA TOP200_DOWN_EVI1 | -1,326 | 0,026 | 0,126 |
| 24 | NOLAN_IFPC | -1,266 | 0,097 | 0,175 |
| 25 | FISCHER_DOWN IN SAA | -1,240 | 0,038 | 0,195 |
| 26 | FISCHER_SAA_RC_DOWN | -1,232 | 0,021 | 0,190 |

| | | | | |
|---|---|---|---|---|
| 27 | SUZ12_DOWN | -1,213 | 0,000 | 0,203 |
| 28 | LAURENTI_CMP | -1,208 | 0,029 | 0,197 |

**ML-DS signature genesets[7]**

| # | Gene set | NES | NOM p | FDR q |
|---|---|---|---|---|
| 1 | ML-DS TOP250 DOWN.GRP | 1,374 | 0,007 | 0,044 |
| 2 | ML-DS TOP400 DOWN.GRP | 1,369 | 0,006 | 0,024 |
| 3 | ML-DS TOP600 DOWN.GRP | 1,352 | 0,000 | 0,020 |
| 4 | ML-DS TOP750 DOWN.GRP | 1,341 | 0,000 | 0,017 |
| 5 | ML-DS TOP500 DOWN.GRP | 1,337 | 0,000 | 0,015 |
| 6 | ML-DS TOP300_DOWN GENE.GRP | 1,319 | 0,022 | 0,017 |
| 7 | ML-DS TOP100_DOWN GENE.GRP | 1,270 | 0,084 | 0,030 |
| 8 | ML-DS TOP400 UP.GRP | -1,734 | 0,000 | 0,000 |
| 9 | ML-DS TOP300_UP GENE.GRP | -1,670 | 0,000 | 0,000 |
| 10 | ML-DS TOP250 UP.GRP | -1,658 | 0,000 | 0,000 |
| 11 | ML-DS TOP500 UP.GRP | -1,642 | 0,000 | 0,000 |
| 12 | ML-DS TOP100_UP GENE.GRP | -1,632 | 0,000 | 0,000 |
| 13 | ML-DS TOP600 UP.GRP | -1,591 | 0,000 | 0,000 |

### Supplemental Figures

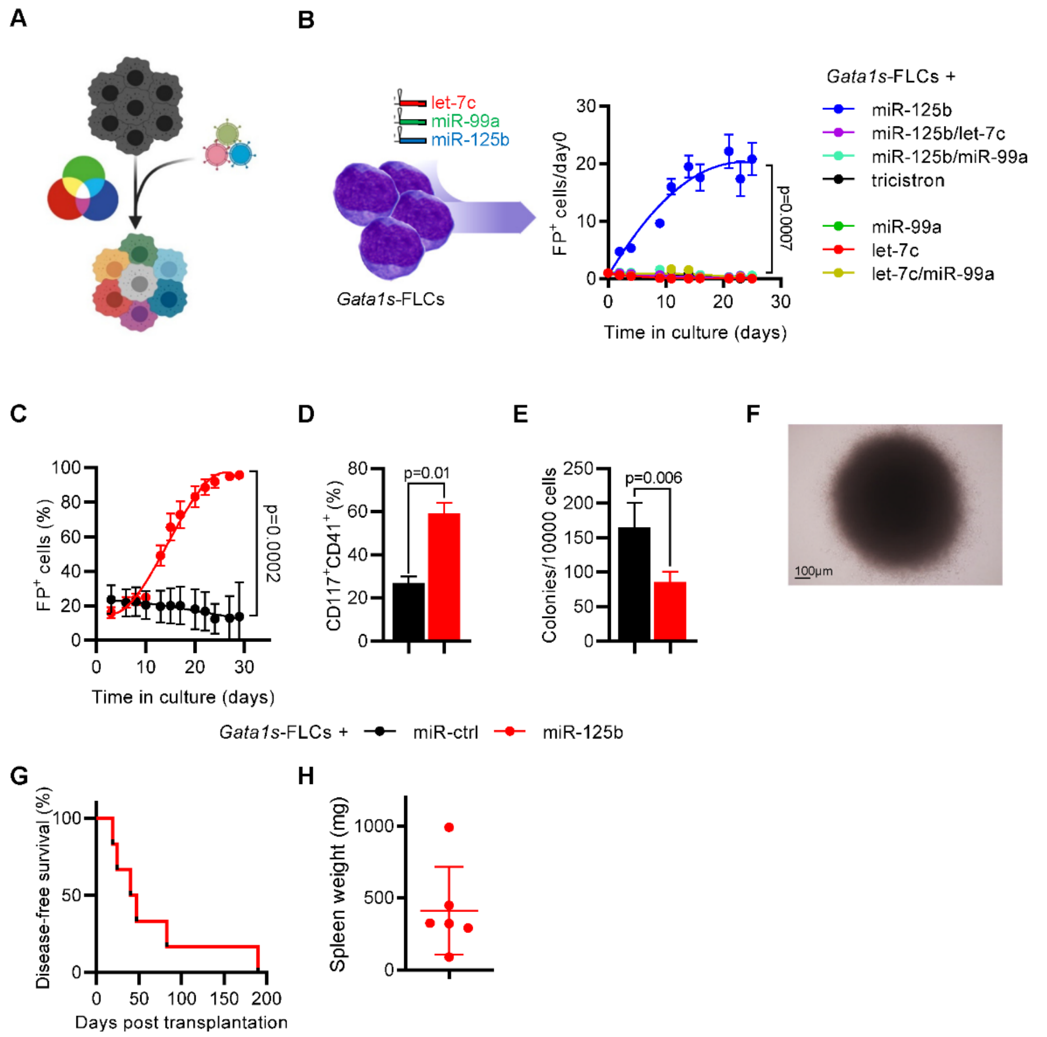


**Supplemental Figure 1. Validation of miR-125b as the only member of the miR-99a~125b tricistrons synergizing with *Gata1s*.**

(A) Schematic of Red-Green-Blue-based lentiviral co-transduction [21]. Co-transduction with different fluorescent protein reporters allows for multicolor tracking and growth competition analysis of independent clonal populations.

(B) Percentage of *Gata1s*-FLCs transduced with different miRNA permutations (marked by dTomato [let-7c], mTagBFP2 [miR-125b] and GFP [miR-99a]) normalized to day 0 (n=4, paired t-test) (right).

(C) Percentage of transduced (miR-ctrl or miR-125b) *Gata1s*-FLCs (n=4, paired t-test).

(D) Bar graph showing the percentage of megakaryocytic progenitors (CD117$^+$CD41$^+$) after 6 days of differentiation. Cells shown are *Gata1s*-FLCs transduced with miR-125b or miR-ctrl (n=3, paired t-test).

E) Bar graph showing the number of colonies generated by transduced *Gata1s*-FLCs (miR-125b or miR-Control) in methylcellulose-based CFU-assays in complete cytokine conditions (n=4).

(F) Representative blast-like colony from one of n=3 independent methylcellulose-based CFU-assays after plating *Gata1s*-FLCs transduced with miR-125b in minimal (Thpo 20ng/ml) cytokine conditions. Scale is indicated.

(G-H) Kaplan-Meier survival curve of secondary recipients transplanted with *Gata1s*-FLCs transduced with miR-125b (25% of bone marrow of primary leukemic recipients) (n=6) and spleen weight upon sacrifice (G).

Data are presented as mean ± SD.

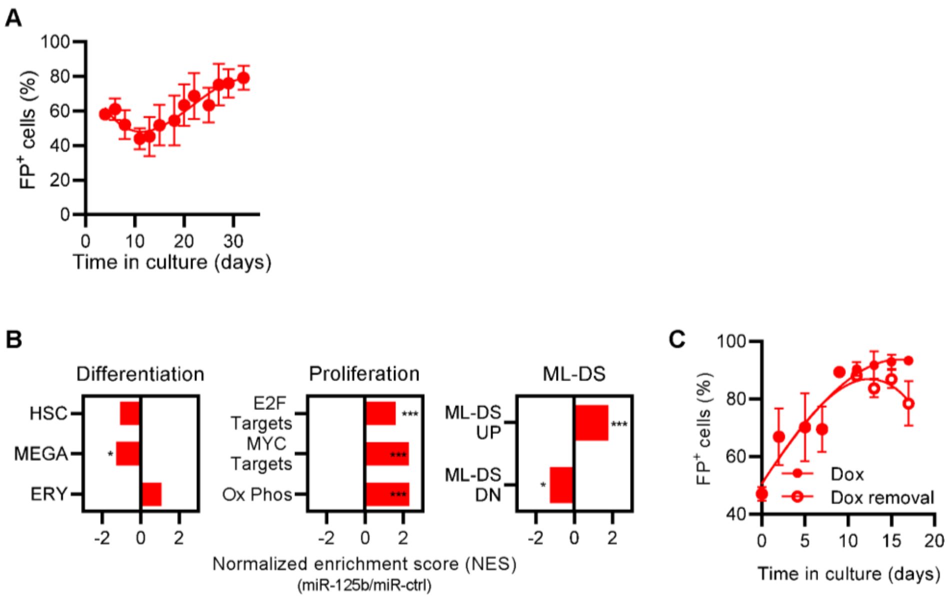


**Supplemental Figure 2. Sustained miR-125b expression induces an ML-DS-like gene expression signature in *Gata1s*-FLCs.**

(A) Percentage of *Gata1s*-FLCs overexpressing the miR-125b-mimic shRNA pool (n=6).

(B) Bar graph showing normalized enrichment scores from GSEA of up- or downregulated gene sets involved in hematopoietic differentiation and cell proliferation after 10 days of doxycycline induction. *Gata1s*-FLCs overexpressing miR-125 were compared to *Gata1s*-FLCs overexpressing miR-ctrl. *= p<0.05; ** = p<0.01; ***=p<0.001.

(C) Percentage of *Gata1s*-FLCs expressing doxycycline-regulated miR-125b upon removal of doxycycline on day 9 of culture (n=2).

All data are presented as mean ± SD.

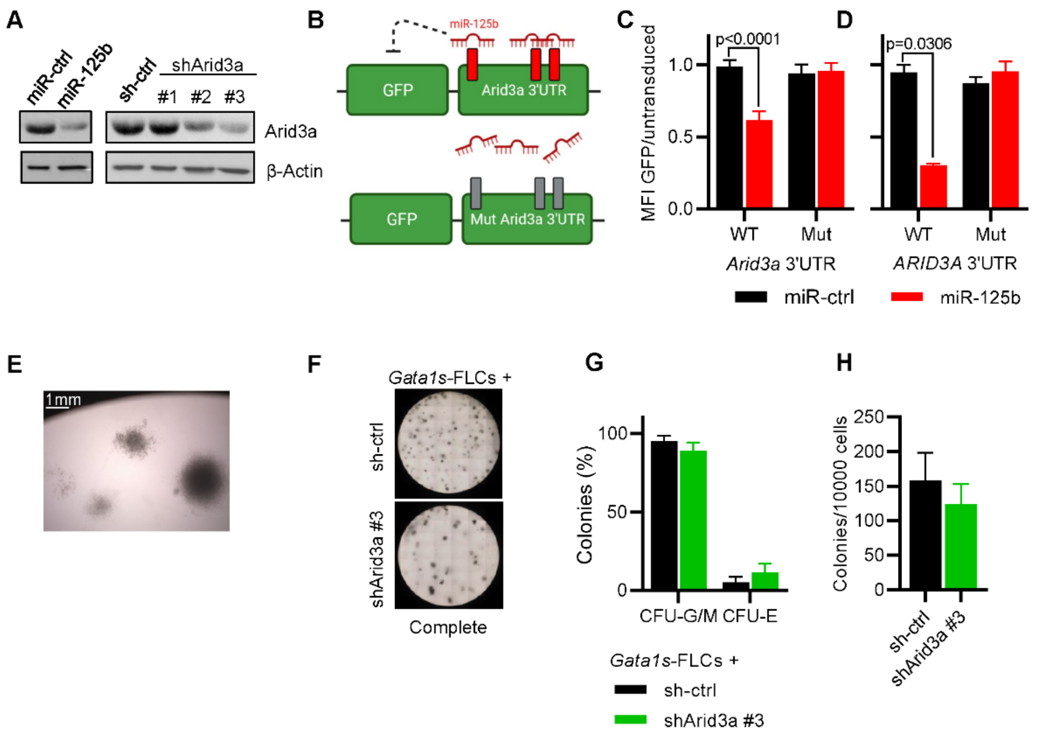


**Supplemental Figure 3. Verification of *Arid3a* as the main target of miR-125b synergizing with *Gata1s*.**

(A) Western Blot showing Arid3a protein levels in *Gata1s*-FLCs 4 days after transduction with miR-125b or shRNAs targeting Arid3a, as well as their respective controls. Position of Arid3a (63kDa) and β-Actin (43kDa) is indicated.

(B) Experimental design of *ARID3A* 3'UTR assays. miR-125b can bind to the 3'UTR and impair GFP expression only when the miR-125b-binding sites are intact.

(C and D) GFP Mean fluorescence intensity of HEL cells expressing wildtype or a mutated 3'UTR of *Arid3a* (B) or *ARID3A* (C) after transduction with miR-125b or miR-Control. MFI is normalized to untransduced cells. (n=12 and n=3, respectively; unpaired t-test).

(E) Representative megakaryocytic-like colony from one of n=3 independent methylcellulose-based CFU-assays after plating *Gata1s*-FLCs transduced with shArid3a #3 in minimal (Thpo 20ng/ml) cytokine conditions.

(F-H) Representative image (F), classification of colonies (G) and number of formed colonies (H) after plating transduced (sh-ctrl (black) or shArid3a #3 (green)) *Gata1s*-FLCs in methylcellulose-based CFU assays under complete cytokine conditions (n=3, paired t-test). CFU-G/M: granulocytic (CFU-G), monocytic (CFU-M) and granulocytic/monocytic (CFU-GFM); CFU-E: erythroid.

All data are presented as mean ± SD.

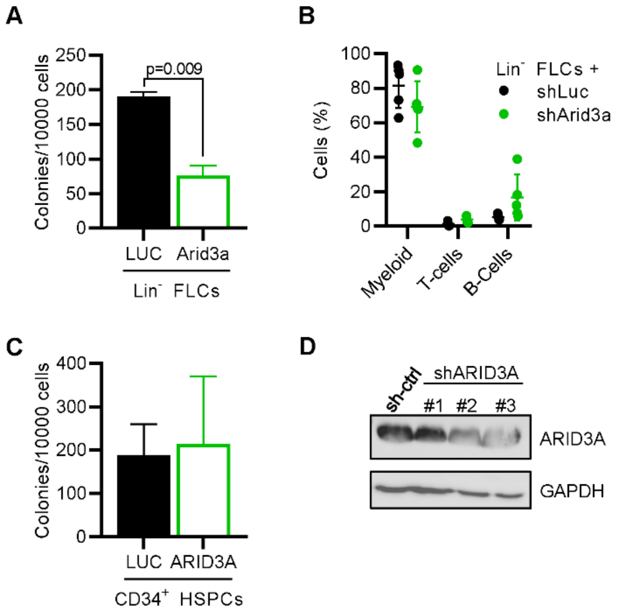


**Supplemental Figure 4. Number of CFUs formed in methylcellulose-based assays and knockdown efficiency of shRNAs against *ARID3A*.**

(A) Number of formed colonies after plating transduced (*Arid3a* or Luc cDNA) murine FLCs in methylcellulose-based CFU assays under complete cytokine conditions (n=2, unpaired t-test).

(B) Bar graph showing the percentage of myeloid ($CD11b^+$), T-cells ($CD3e^+$) and B-cells ($B220^+$) in the BM of mice transplanted with $Lin^-$ FLCs transduced with sh-ctrl (black) or shArid3a (green). $shRNA^+$ cells shown. n=5.

(C) Number of formed colonies after plating *ARID3A*- or Luc-expressing human $CD34^+$ HSPCs in methylcellulose-based CFU assays under complete cytokine conditions (n=3, paired t-test).

(D) Western Blot showing ARID3A protein levels in K562 4 days after transduction with shRNAs targeting *ARID3A*, as well as a non-targeting control (sh-ctrl). Position of ARID3A (63kDa) and GAPDH (37kDa) is indicated.

All data are presented as mean ± SD.

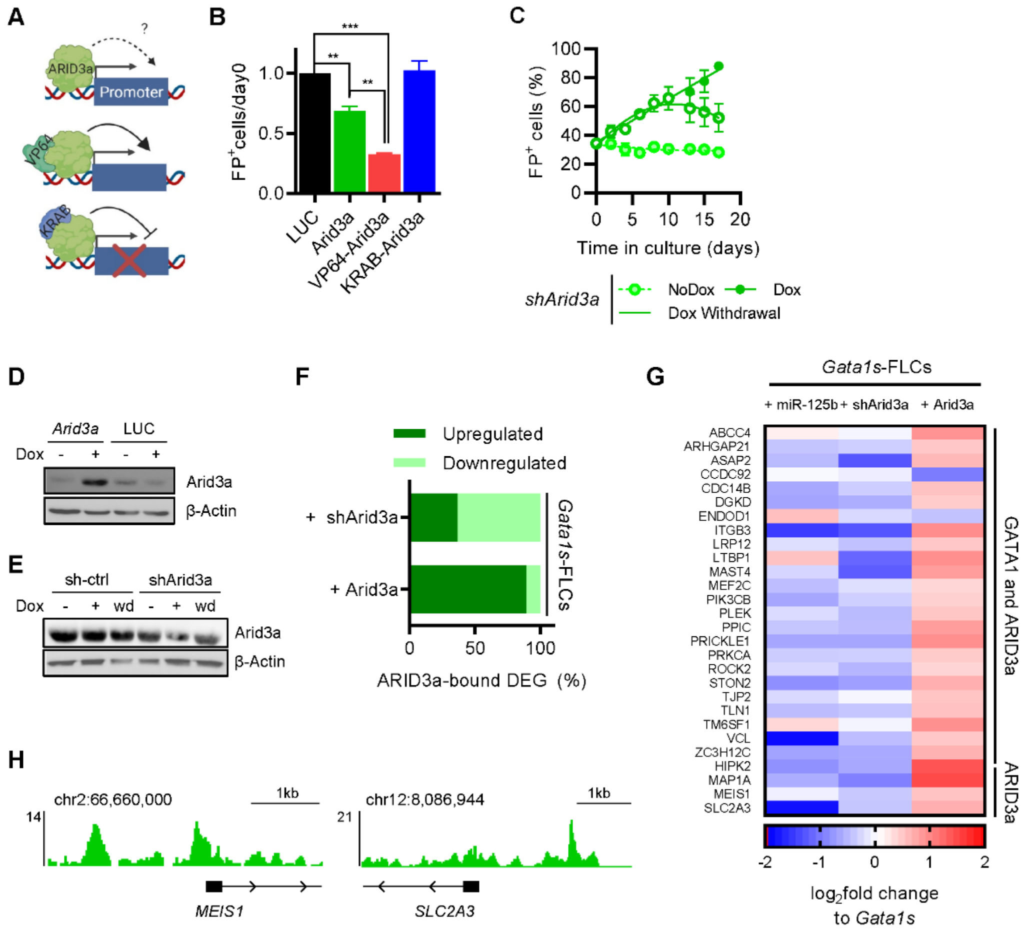


**Supplemental Figure 5. ARID3A activates expression of genes involved in megakaryocytic differentiation.**

(A-B) Schematic of setup to determine the role of ARID3A in gene transcription by fusing the VP64-activator or KRAB-inhibitor domains to its N-terminal domain (A) and normalized percentage (to LUC cDNA) of transduced (ARID3A (green), VP64-ARID3A (red) or KRAB-ARID3A (blue)) *Gata1s*-FLCs after 10 days in culture. n=3; *= p<0.05; ** = p<0.01; ***=p<0.001 (paired t-test).

(C and D) Western Blot showing Arid3a protein levels of *Gata1s*-FLCs 4 days after doxycycline-mediated induction of *Arid3a* (C) or shArid3a (D) expression (or its respective controls). Samples after doxycycline withdrawal (wd) were taken 2 days after removal of doxycycline. Position of Arid3a (63kDa) and β-Actin (43kDa) is indicated.

(E) Percentage of *Gata1s*-FLCs expressing doxycycline-regulated shArid3a upon addition or removal of doxycycline (500ng/mL; n=3).

(F) Expression profile of differentially expressed genes bound by ARID3A in *Gata1s*-FLCs after induction of *Arid3a* or shArid3a (n=157 and n=490, respectively).

(G) Gene expression profile of genes involved in megakaryocytic differentiation upon modulation of *Arid3a* expression in *Gata1s*-FLCs. Data shown as $log_2$fold change to each respective control, binding by ARID3A or GATA1 and ARID3A is indicated (right).

(H) IGV snapshot showing occupancy of megakaryocytic genes by ARID3A (K562, ENCODE datasets).

Data are presented as mean ± SD.

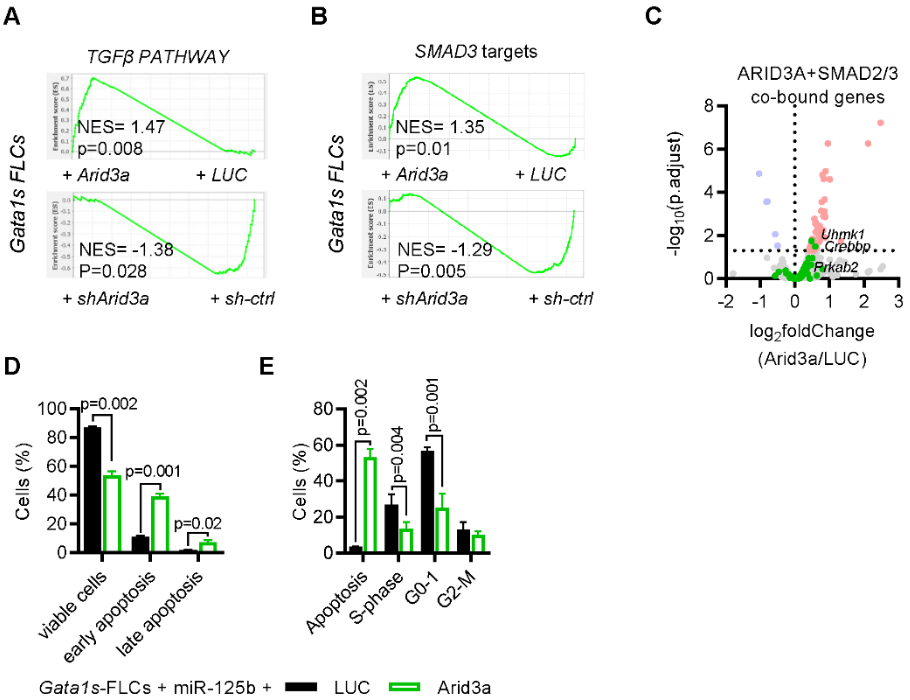


**Supplemental Figure 6. ARID3A binds to SMAD2/3 activating TGFβ-mediated apoptosis and cell cycle arrest.**

(A-B) GSEA enrichment plots showing the modulation of the TGFβ signaling pathway (A) and induction of SMAD3 targeted genes (B) upon *Arid3a* induction or repression in *Gata1s*-FLCs.

(C) Volcano plot showing expression of genes bound by the ARID3A-SMAD2/3 complex upon induction of *Arid3a* expression in *Gata1s*-FLCs. Genes involved in apoptosis and cell cycle arrest are highlighted in green; significantly downregulated and upregulated genes in blue and red, respectively.

(D) Percentage of viable, early and late apoptotic *Gata1s*-FLCs overexpressing miR-125b after 7 days of doxycycline induction of *ARID3A* expression as determined by Annexin-V assay. Viable cells: Annexin-V$^-$/DAPI$^-$; early apoptotic: Annexin-V$^+$/DAPI$^-$; late apoptotic: Annexin-V$^+$/DAPI$^+$. n=3 (paired t-test).

(E) Percentage of *Gata1s*-FLCs overexpressing miR-125b in apoptosis, S-phase, G0-1 and G2-M after 7 days of doxycycline induction of *ARID3A* expression as determined in BrDU-based cell cycle assay. Apoptosis: BrdU$^-$/7-AAD$^-$; G0-1: BrdU$^-$/7-AAD $^{low}$; S-Phase: BrdU$^+$/7-AAD $^+$; G2-M: BrdU$^-$/7-AAD $^{high}$. n=3 (paired t-test).

Data are presented as mean ± SD.

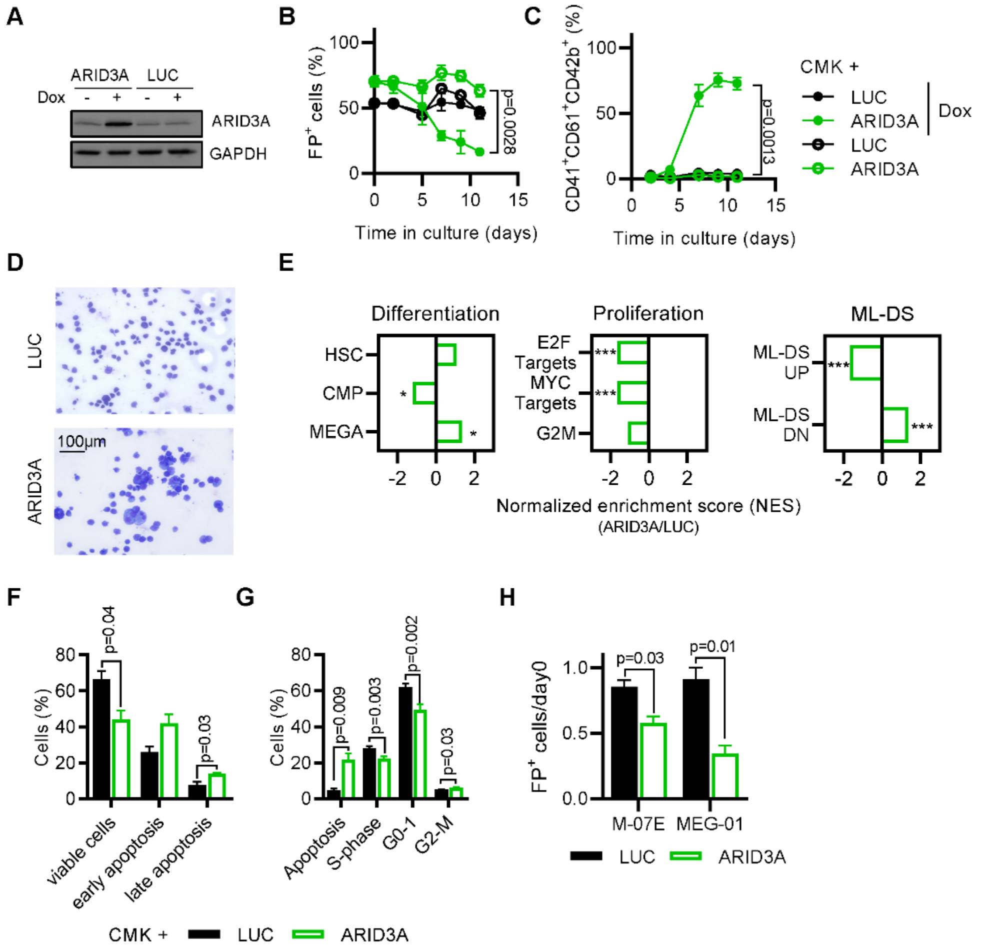


**Supplemental Figure 7. Overexpression of *ARID3A* in the ML-DS cell line CMK restores differentiation potential and mediates cell cycle arrest and apoptosis.**

(A) Western Blot showing ARID3A protein levels in CMK cells 4 days after doxycycline-mediated induction of *ARID3A* or LUC cDNA expression. Position of ARID3A (63kDa) and GAPDH (37kDa) is indicated.

(B-D) Percentage of fluorescent reporter positive (A) and percentage of mature megakaryocytic cells ($CD41^{+}CD61^{+}CD42b^{+}$) (B) after doxycycline induction of CMK cells transduced with dox-inducible *ARID3A* or LUC cDNAs. (C) Representative micrograph of n=3 experiments (paired t-test).

(E) Bar graphs showing normalized enrichment scores from GSEA of up- or downregulated gene sets involved in hematopoietic differentiation, cell proliferation and ML-DS progression. CMK overexpressing *ARID3A* were compared to CMK overexpressing LUC cDNA two days after doxycycline induction. *= p<0.05; ** = p<0.01; ***=p<0.001.

(F) Percentage of viable, early and late apoptotic CMK cells after 7 days of doxycycline induction of *ARID3A* expression as determined by Annexin-V assay. Viable cells: Annexin-$V^{-}$/$DAPI^{-}$; early apoptotic: Annexin-$V^{+}$/$DAPI^{-}$; late apoptotic: Annexin-$V^{+}$/$DAPI^{+}$. n=3 (paired t-test).

(G) Percentage of CMK cells in apoptosis, S-phase, G0-1 and G2-M after 7 days of doxycycline induction of *ARID3A* expression as determined in BrDU-based cell cycle assay. Apoptosis: $BrdU^{-}$/7-$AAD^{-}$; G0-1: $BrdU^{-}$/7-$AAD^{low}$; S-Phase: $BrdU^{+}$/7-$AAD^{+}$; G2-M: $BrdU^{-}$/7-$AAD^{high}$. n=3 (paired t-test).

(H) Normalized (to LUC) percentage of M-07E or MEG-01 cells expressing constitutively expressed *ARID3A* after 10 days in culture (n=3, paired t-test).

All data are presented as mean ± SD.

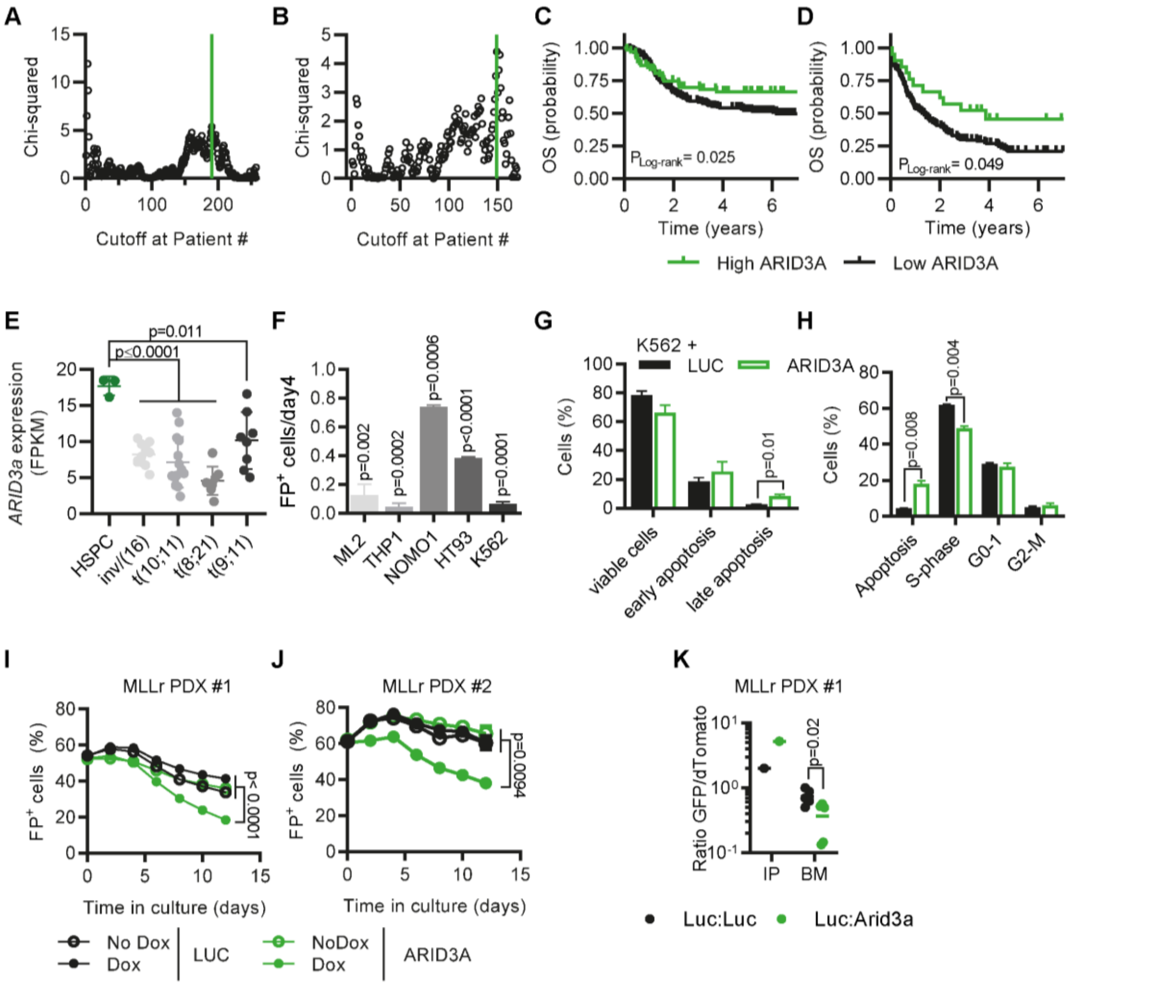


**Supplemental Figure 8. ARID3A is a global tumor suppressor in AML**

(A-B) Event-free survival (EFS) statistic (Chi-squared) as function of the cutoff point in the NCI-TARGET (A) and the TCGA (B) patient populations. The calculated optimum cutoff for EFS (green line, as determined by maximally selected rank statistics) – 12.0 and 12.3 normalized ARID3a reads, respectively – was used for both OS and EFS computations.

(C) Probability of overall survival in 258 pediatric AML patients with high (green; >12.0 normalized reads) or low ARID3a expression (black; ≤12.0 normalized reads).

(D) Probability of overall survival in 171 adult AML patients with high (green; >12.3 normalized reads) or low ARID3a expression (black; ≤12.3 normalized reads).

(E) *ARID3A* expression (FPKM) in sorted pediatric AML blasts of different non-AMKL subtypes and $CD34^+$ HSPCs.

(F) Bar graph showing the percentage of $ARID3A^+$ cells after 7 days of induction with doxycycline, normalized to the LUC control.

(G) Percentage of viable, early and late apoptotic K562 cells after 7 days of doxycycline induction of *ARID3A* expression as determined by Annexin-V assay. Viable cells: Annexin-$V^-$/$DAPI^-$; early apoptotic: Annexin-$V^+$/$DAPI^-$; late apoptotic: Annexin-$V^+$/$DAPI^+$. n=3 (paired t-test).

(H) Percentage of K562 cells in apoptosis, S-phase, G0-1 and G2-M after 7 days of doxycycline induction of *ARID3A* expression as determined in BrDU-based cell cycle assay. Apoptosis: $BrdU^-$/7-$AAD^-$; G0-1: $BrdU^-$/7-$AAD^{low}$; S-Phase: $BrdU^+$/7-$AAD^+$; G2-M: $BrdU^-$/7-$AAD^{high}$. n=3 (paired t-test).

(I-J) MLLr PDXs #1 and #2 were transduced with doxycycline-inducible *ARID3A* or LUC cDNA vectors. Percentage of fluorescent reporter positive cells after doxycycline induction of MLLr PDX #1 (I) or MLLr PDX #2 (J) (n=3, paired t-test)

(K) Ratio of $GFP^+$ to $dTomato^+$ cells in input cells (IP), and in the BM of mice sacrificed 4-5 weeks after transplantation of MLLr PDX #1 transduced with *ARID3A* ($GFP^+$) or a LUC control ($GFP^+$) and mixed 1:1 with LUC control-transduced blasts ($dTomato^+$) before transplantation (n=5, unpaired t-test).

### Supplemental Information References

1. Gialesaki S, Mahnken AK, Schmid L, et al. GATA1s exerts developmental stage-specific effects in human hematopoiesis. *Haematologica*. 2018;103(8):e336-e340.
2. Labuhn M, Perkins K, Matzk S, et al. Mechanisms of Progression of Myeloid Preleukemia to Transformed Myeloid Leukemia in Children with Down Syndrome. *Cancer Cell*. 2019;36(2):123-138.e110.
3. Emmrich S, Rasche M, Schöning J, et al. miR-99a/100~125b tricistrons regulate hematopoietic stem and progenitor cell homeostasis by shifting the balance between TGFβ and Wnt signaling. *Genes & development*. 2014;28(8):858-874.
4. Klusmann JH, Li Z, Böhmer K, et al. miR-125b-2 is a potential oncomiR on human chromosome 21 in megakaryoblastic leukemia. *Genes Dev*. 2010;24(5):478-490.
5. Adams FF, Heckl D, Hoffmann T, et al. An optimized lentiviral vector system for conditional RNAi and efficient cloning of microRNA embedded short hairpin RNA libraries. *Biomaterials*. 2017;139:102-115.
6. Fellmann C, Zuber J, McJunkin K, et al. Functional identification of optimized RNAi triggers using a massively parallel sensor assay. *Mol Cell*. 2011;41(6):733-746.
7. Schwarzer A, Emmrich S, Schmidt F, et al. The non-coding RNA landscape of human hematopoiesis and leukemia. *Nat Commun*. 2017;8(1):218.
8. Wiśniewski JR, Zougman A, Nagaraj N, Mann M. Universal sample preparation method for proteome analysis. *Nature Methods*. 2009;6(5):359-362.
9. Consortium TU. UniProt: the universal protein knowledgebase in 2021. *Nucleic Acids Research*. 2020;49(D1):D480-D489.
10. Eng JK, McCormack AL, Yates JR. An approach to correlate tandem mass spectral data of peptides with amino acid sequences in a protein database. *Journal of the American Society for Mass Spectrometry*. 1994;5(11):976-989.
11. Labuhn M, Adams FF, Ng M, et al. Refined sgRNA efficacy prediction improves large- and small-scale CRISPR-Cas9 applications. *Nucleic Acids Res*. 2018;46(3):1375-1385.
12. Chen S, Zhou Y, Chen Y, Gu J. fastp: an ultra-fast all-in-one FASTQ preprocessor. *Bioinformatics*. 2018;34(17):i884-i890.
13. Spinozzi G, Tini V, Mincarelli L, Falini B, Martelli M. A comprehensive RNA-Seq pipeline includes meta-analysis, interactivity and automatic reporting; 2018.
14. Love MI, Huber W, Anders S. Moderated estimation of fold change and dispersion for RNA-seq data with DESeq2. *Genome Biology*. 2014;15(12):550.
15. Benjamini Y, Hochberg Y. Controlling the False Discovery Rate: A Practical and Powerful Approach to Multiple Testing. *Journal of the Royal Statistical Society: Series B (Methodological)*. 1995;57(1):289-300.
16. Subramanian A, Tamayo P, Mootha VK, et al. Gene set enrichment analysis: a knowledge-based approach for interpreting genome-wide expression profiles. *Proc Natl Acad Sci U S A*. 2005;102(43):15545-15550.
17. Aken BL, Achuthan P, Akanni W, et al. Ensembl 2017. *Nucleic Acids Res*. 2017;45(D1):D635-d642.
18. Kaplan EL, Meier P. Nonparametric Estimation from Incomplete Observations. *Journal of the American Statistical Association*. 1958;53(282):457-481.
19. Mantel N. Evaluation of survival data and two new rank order statistics arising in its consideration. *Cancer Chemother Rep*. 1966;50(3):163-170.
20. Leek JT, Johnson WE, Parker HS, Jaffe AE, Storey JD. The sva package for removing batch effects and other unwanted variation in high-throughput experiments. *Bioinformatics*. 2012;28(6):882-883.
21. Weber K, Thomaschewski M, Warlich M, et al. RGB marking facilitates multicolor clonal cell tracking. *Nat Med*. 2011;17(4):504-509.

22. Liberzon A, Birger C, Thorvaldsdóttir H, Ghandi M, Mesirov JP, Tamayo P. The Molecular Signatures Database (MSigDB) hallmark gene set collection. *Cell systems*. 2015;1(6):417-425.
